# Negative Mem-Capacitance and Warburg Ionic Filtering in Asymmetric Nanopores

**DOI:** 10.1101/2022.10.20.513121

**Authors:** Nasim Farajpour, Y. M. Nuwan, D. Y. Bandara, Vinay Sharma, Lauren Lastra, Kevin J. Freedman

## Abstract

The pervasive model for a solvated, ion-filled nanopore is often a resistor in parallel with a capacitor. However, for conical nanopore geometries, we propose the inclusion of a Warburg-like element which is necessary to explain otherwise anomalous observations such as negative capacitance and lowpass filtering of translocation events (*i.e.,* a phenomenon we term Warburg filtering). The negative capacitance observed here is characterized as having long equilibration times and memory (*i.e.,* mem-capacitance) at negative voltages. Next, we used the transient occlusion of the pore using λ-DNA and 10-kbp DNA to test whether events are being attenuated by purely ionic phenomena even when there is sufficient amplifier bandwidth. The inclusion of the Warburg-like element is mechanistically linked to concentration polarization and the activation energy to generate and maintain localized concentration gradients. We conclude the study with a new interpretation of molecular translocations which is not simply based on the pulse-like resistance changes but rather a complex and non-linear storage of ions that changes during molecular transit.

## Introduction

Principles of non-linear electrokinetics are often used to explain otherwise anomalous data produced by electrochemical cells^1^, ionic liquids^2^, nanofluidic devices^3,4^, and other systems^5–7^. The classical Poisson-Nernst-Planck and Navier-Stokes equations represent a useful approximation of current and ionic concentrations however tends to deviate from experiments under (1) crowded conditions and (2) strong surface interactions^8–10^. Although these two conditions describe a nanopore filled with an electrolyte, the impact of non-linear electrokinetics on single molecule sensing experiments has been sparsely explored. One of the dominant mechanisms linked to non-linear phenomenon is non-linear capacitance stemming from ion interactions with a charged surface. Distinct from membrane (e.g. solid-state) capacitance, the electric double layer (EDL), which is composed of both Stern and diffuse layers and described by the Gouy-Chapman-Stern (GCS) model^11^, also contributes its own capacitance. More importantly, the EDL capacitance is ion-specific and concentration-dependent which are routinely varied within nanopore experiments^8,12,13^. Since capacitive elements are capable of storing and releasing charge, they are important to consider when making ultra-small (*i.e.,* pico-ampere) current measurements at nanoscale interfaces. Nanopipettes, in particular, are well-suited for studying non-linear ionic phenomenon since the glass walls are highly charged and the geometry confines ions and hinders diffusion to virtually one axis which slows down charge migration^14^.

Electrical circuit equivalents, such as resistive and capacitive (RC) components, have routinely been used to model ideal charge transport in ionic systems^15,16^. The nanopore field has utilized these models to help understand the concepts of access resistance^17^, sensing zone^18^, capture radius^19^, and capacitive noise^20,21^; the latter of which has been the theoretical basis for low noise nanopore systems^22^. While the resistive nature of a nanopore is undisputed, the capacitive components of the nanopore system are undoubtedly more complex; some capacitive phenomenon (e.g. “mem-capacitance”) are only revealed through simulations^23^. Broadly, capacitive current can fall under two categories: (1) leaky capacitance wherein the voltage is not changing 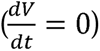 yet current can still traverse the capacitor, and (2) the more canonical capacitance wherein current only flows when there is a voltage change 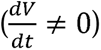. Leaky capacitance will be discussed here since the EDL can physically store charge as well as conduct charge along interfacial surfaces^24^. A leaky capacitor can be modeled as a parallel RC circuit in which charge is both stored as well as passes a steady-state current. Conceptually, since the EDL can contribute to the conduction of ions through the nanopore, a leaky capacitor model may be fitting in some situations, such as low (mM) salt conditions. In both leaky and canonical capacitance, negative capacitance is also possible which refers to a system that stores less charge (Q) with increasing voltage (V), has been reported in ionic systems^23,25,26^. The characteristic feature of negative capacitance is that the equilibrium (*i.e.,* resistive) current is higher than the combined current (*i.e.,* resistive and capacitive combined).

Planar and conical solid-state nanopores are routinely used for single molecule sensing however ionic phenomenon such as concentration polarization, rectification, electroosmotic flow reversal, and negative resistance have only been observed within nanopipettes. Concentration polarization is observed as a reduction or enhancement of the conductivity of the fluid inside the tapered region of the nanopipette^27–29^. Under low frequency voltage stimulation or voltage steps, the formation of a concentration gradient is governed by diffusional transport and thus associated with a time constant. Motivated by the models typically used in electrochemical impedance spectroscopy, diffusion-limited processes (namely, semi-infinite linear diffusion) can be described by a Warburg element that has an associated Warburg impedance (Z_w_)^30–33^. The equivalent circuit for a Warburg element is several RC elements arranged in series to account for multiple time constants^34–37^. The relationship between a Warburg element and a leaky capacitor is also noteworthy since both can be modeled as RC elements. The region where the Warburg element is most likely to apply to nanopipettes is the tapered region wherein there is roughly one-dimensional diffusion of ions. Since the Warburg element is in series with the nanopore, the impact on current sensing would be significant and currently has not been studied to our knowledge.

In this study, investigate the impact of placing a simplified Warburg element (a single RC element) in series with a current-limiting resistor (*i.e.,* the resistance associated with the nanopore) and highlight the significance it plays in ionic current sensing. Three ionic phenomenon are studied in particular: negative capacitance, mem-capacitance, and ionic filtering (an effect we call Warburg filtering). Warburg filtering refers to the poor ability to resolve fast transient events due to a lowpass filtering effect that is distinct from instrument lowpass filtering. With the recent interest in high-bandwidth instrumentation for nanopore measurements, understanding the fundamental ionic limits of conical nanopore recordings is important for future advances. Throughout this study, capacitance effects in conical nanopores and the circuit architecture itself will be explored. In addition to low salt conditions (10 mM KCl), salt gradients were included here due to several reports of non-linear electrokinetic behavior^1,8,38,39^. Both current-voltage (IV) data and translocation data will be used to explore the nature of capacitance in nanopipettes and how the capacitance may influence the collection of single molecule data.

## Results and Discussion

### Circuit Model of a Conical Nanopore

In this work, we propose that a Warburg element plays a critical role in ionic current measurements within conical nanopores. Modeling conical nanopores with an RC element in series with a resistor (*i.e.,* the nanopore) has significant implications for understanding current fluctuations during a voltage switch as well as during molecular transit. Since the Warburg elements can be simplified into an RC component, a new circuit model is proposed, as shown in **Figure 1a**. The rationale for including the Warburg element in the equivalent circuit is based on the need for an associated time constant for which concentration polarization can reach equilibrium (*i.e.,* concentrations do not change instantaneously). The capacitor, C_W_, therefore can be thought of as the activation energy required to generate and maintain concentration polarization. Furthermore, charge migration will be limited by diffusion within the confinement of the nanopipette. In the analysis to follow, the circuit can further be analyzed under steady state (*i.e.,* voltage is constant and current has equilibrated) and dynamic states (*i.e.,* voltage is not constant). Under steady state conditions, only the resistors would play a role in limiting current, and given the high-aspect ratio of nanopipettes, it is reasonable to represent the tapered region by its own resistor which is in series with pore resistance. Furthermore, the resistance is given by: 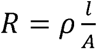, where l, A, and ρ are the length, cross-sectional area, and resistivity of the fluid, respectively. Therefore, at steady-state (ignoring capacitive elements), the equivalent circuit is two resistors in series. At positive voltages, the resistivity of the electrolyte inside the taper is reduced causing the total resistance (R_W_+R_pore_) to increase; this is typically seen as rectification of the ionic current at the positive polarity (**see Supplemental Information**). A finite element model was used to demonstrate the concentration polarization phenomenon within a 30 nm pore filled with 10 mM KCl at -1000 mV and +1000 mV (**Fig. 1b**, see **Supplemental Information** for additional details). At +1000 mV, the pore (*i.e.,* smallest constriction) had an average concentration of 9.5 mM, whereas 100 nm into the nanopipette had an average concentration of 3.5 mM (*i.e.,* ion depleted state). At -1000 mV, the pore had an average concentration of 9.6 mM, whereas 100 nm into the nanopipette had an average concentration of 16.9 mM (*i.e.,* ion enriched state).

**Fig 1.**
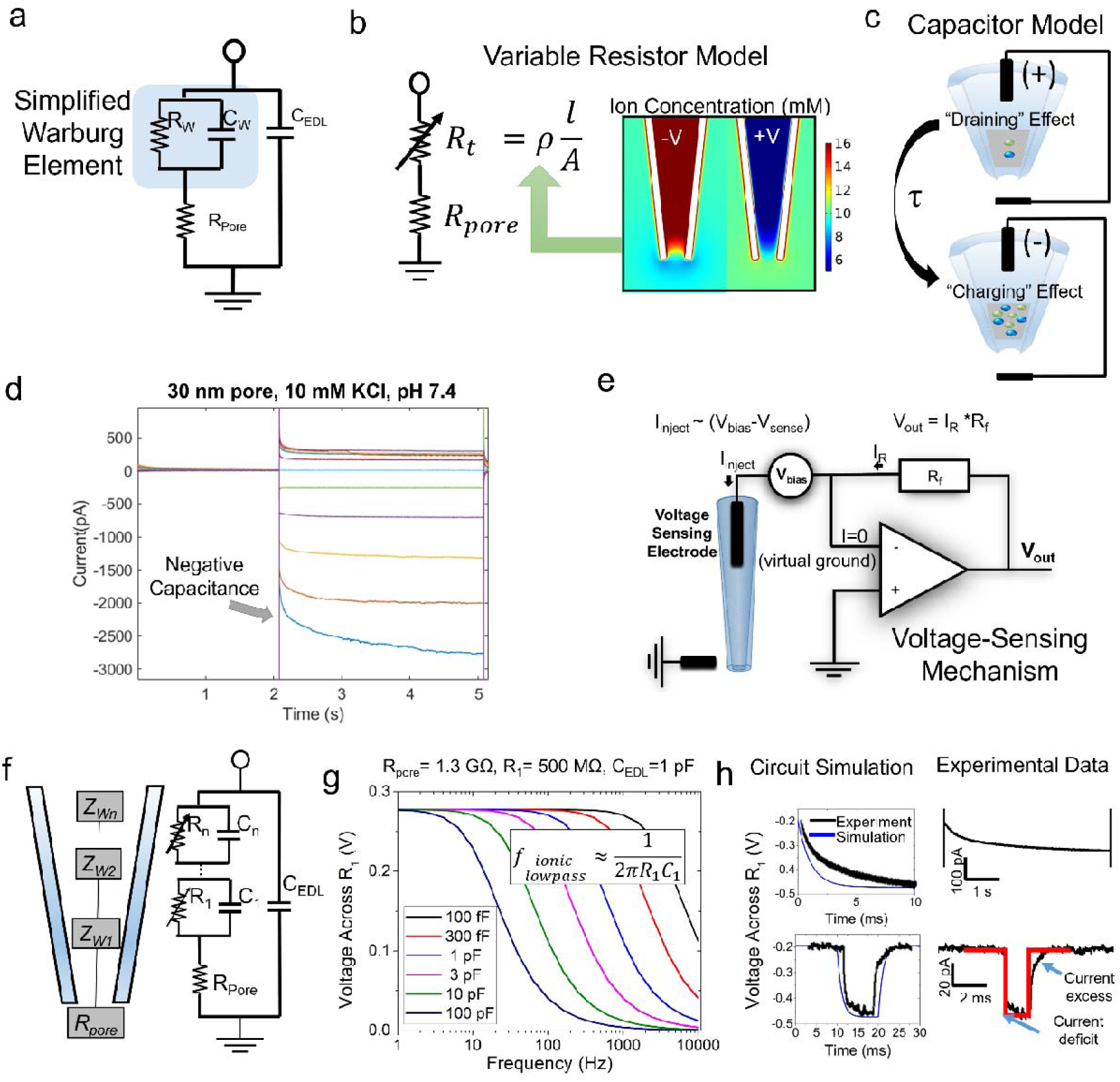
(a) Equivalent circuit model for conical nanopores which include an RC element corresponding to the tapered region. (b) Steady-state circuit equivalent with two distinct resistors representing the pore and the taper. The resistor representing the tapered region i shown as being a variable resistor due to the effects of concentration polarization. Positive voltages deplete or “drain” the taper of ions, while negative voltages enrich or charge the taper with ions. (c) Dynamically changing voltage conditions lead a portion of the nanopore ionic flux to be stored or released from the tapered region resulting in charging/discharging behavior. (d) Experimental data showing the raw data of sequential voltage pulses used to generate a current-voltage curve. The pore diameter is approximated to be ∼30 nm and was filled with 10 mM KCl (pH 7.4). (e) Schematic of a typical transimpedance amplifier which injects and senses current based on the voltage sensed at the electrode inside the nanopipette. (f) The proposed equivalent circuit model for a nanopipette which contains a Warburg element in series with the nanopore resistance. The Warburg element is modeled as an infinite series of RC elements. (g) Frequency domain simulation of the voltage across R1 which represents a Warburg element with n=1. The current through R_1_ also exhibits the same attenuation properties as the voltage across R_1_. (h) Comparative analysis of experimental data and the circuit simulation results for a voltage step and transient pulse. The voltage step was acquired from a ∼30 nm pore symmetrically filled with 10 mM KCl (pH 7.4) and after -1000 mV was applied to the pore (zero voltage bias prior to the voltage pulse). The transient pulse was obtained by translocating λ-DNA through a ∼20 nm pore at 10 mM KCl (V=+500 mV, pH 7.4). Circuit simulations were performed using CircuitLab.

Switching between the enriched and depleted state is also limited by mass transport (*i.e.,* Poisson-Nernst-Planck) and governed by a diffusion-limited time constant, τ (**Fig. 1c**). Although positive and negative ions are “stored” similar to a conventional capacitor, it should be noted that the tapered region of the nanopipette is a non-standard capacitor wherein charge is co-localized rather than separated in space (*e.g.,* commonly idealized by parallel plates). Despite being co-localized, when a voltage switch occurs, the ions travel in opposing directions similar to the conventional capacitor. Since a portion of ionic flux will be stored in the tapered region of the nanopipette, the observation of “negative capacitance” is expected when switching to a more enriched state (*e.g.,* positive voltage to negative voltage). Conversely, when the voltage bias is changed resulting in a depleted state, excess ions will be transported through the pore leading to positive capacitance or the traditional current spike. Therefore, negative capacitance should only appear when switching the voltage to a negatively charged bias.

Experiments were performed to test the polarity specific observations of negative capacitance; namely a pore (∼30 nm) was filled with 10 mM KCl (pH 7.4) and voltage pulses were applied from -500 mV to +500 mV. Given the equivalent circuit for the nanopipette (**Fig. 1a**), current will only be sensed if ions travel through both R_pore_ and R_W_ which is notably different than the traditional notion that ions are sensed immediately after transiting the pore (R_pore_). Our experiments show that for a voltage step in the negative direction (a negative voltage applied inside the nanopipette), the current gradually increases to a higher conduction state (**Fig. 1d**). This observation can be interpreted as ionic current gradually charging the capacitor, C_t_, until the ionic concentrations for the enriched state reach equilibrium. Unlike positive capacitance which leads to a spike in current, negative capacitance is observed as a transient reduction in current in the moments after a voltage switch. Negative capacitance is most clearly observed in the voltage step from 0 mV to -500 mV (**Fig. 1d**, bottom trace) and is associated with a slow time constant (*i.e.,* the current takes several seconds to reach steady-state). Negative capacitance is also observed for voltage pulses -400, -300, and -200 mV but equilibrate to steady state with a faster time constant. For a voltage step in the positive direction, ionic concentration reduces inside the taper and so ionic current initially is high and decays to a lower state. The resultant current traces mimic the expected positive capacitance that occurs at high electrolyte concentrations (e.g., 1 M KCl) in both planar and conical nanopores^20,40^.

The traditional conceptualization of current sensing through a nanopore seems to be complicated by capacitance; notably when it is in series with the nanopore. Here, we proposed that only current flowing through the tapered region of the nanopipette is measured. The mechanism of current sensing can further be modified by interpreting the current signal as the current required to maintain the voltage between the two electrodes in the system; often referred to as the clamping voltage (V_clamp_). Current is injected, as-needed, to keep the voltage bias from drifting away from the set value (**Fig. 1e**)^41,42^. The voltage across R_W_ therefore becomes a critical factor in understanding the current measured through the nanopore. Distinguishing between current and voltage is only important when there is a possibility of charge accumulation that lead to violations of electroneutrality. In cases where cations and anions are perfectly matched, the voltage distribution is solely dictated by the pore geometry. However, based on Poisson’s equation, the accumulation anions or cations within the nanopipette will generate its own voltage and therefore influence the distribution of voltage within nanopipette. The voltage distribution within the tapered region of the nanopipette is therefore critical to current sensing and is the basis for understanding transient ionic phenomenon within nanopipettes. Specifically, the time constant for charge migration within the tapered region can lead to RC-like transients.

The equivalent circuit model of the nanopipette should be represented by numerous RC elements which would reflect the conical geometry of the nanopipette, as well as more accurately model a theoretical Warburg element (**Fig. 1f**). The implication of having multiple RC elements is that there would be multiple RC time constants which would influence the capacitive response of the nanopore. Indeed, experimentally, the negative capacitance is poorly fitted by a single exponential and is best modelled as a multitude of exponentials with different time constants. For simplicity, only a single RC element (*i.e.,* n=1) will be used for the rest of this study and is suitable for modeling the dominant RC element. It should also be noted that a Warburg element with n=1 is equivalent to a leaky capacitor model. A circuit model was built using the following parameters: R_pore_=1.3 GOhm, R_W_=R_1_=500 MOhm, C_EDL_= 1pF (1.3 GOhm was the resistance of a 30 nm pore in 10 mM KCl). The capacitance of the taper was varied between 100 fF ad 100 pF, and the voltage across R_1_ was plotted as a function of frequency (**Fig. 1g**). The RC element is shown to have a lowpass filtering effect on the current and voltage associated with R_1_. The cutoff frequency of the lowpass filter is dependent on the values of R_1_ and C_1_ and given by: 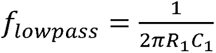.

Time domain circuit simulations (*i.e.,* the circuit shown in **Figure 1f** and n=1) were also performed for R_pore_=1.3 GOhm, R_W_=R_1_=500 MOhm, C_EDL_= 1pF, and C_1_=100pF. For a voltage step from 0V to -500 mV, the transient current response is shown for both the circuit simulation as well as a 30 nm pore (10 mM KCl, pH 7.4). The main difference between experiments and the simulation is that experimental data shows dynamic (*i.e,* decaying) time constant which causes a slower decay towards the equilibrium current (**Fig. 1h**). A voltage pulse was also used to simulate a translocation event. Although the applied voltage pulse was “square”, the capacitor C1 acts as a filter to the high frequency components of the current leading to an RC time constant-associated rise/fall time. At the low salt conditions used here (10 mM KCl, pH 7.4), only conductive events for λ-DNA were observed but with a distinctive exponential decay toward the blocked state and also back to baseline (**Fig. 1h**: V= -500 mV, **see Methods** for experimental details). Interestingly, the current response for a translocation event shows signs of negative capacitance wherein there is a current deficit and current excess at the beginning and end of the event, respectively. Capacitive currents therefore show up in patch-clamp measurements as having a negative contribution to total current. The un-intuitive current responses shown here therefore seems to be the result of: (1) a capacitive element being in series with the nanopore’s resistance, and (2) the inability of voltage-clamp experiments to measure total current, but rather only the current that affects the voltage at the electrode-solution interface.

### Experimental Negative Mem-Capacitance

Nanopipettes were fabricated from laser heating quartz capillaries and pulling them to nanoscale conical tips. Transmission electron microscopy was used to image the tips and obtained their internal diameters as a function of pulling parameters (see **Supplemental Information**). Once fabricated, the pores were filled with electrolyte and Ag/AgCl electrodes were placed inside the nanopipette (V_clamp_) and inside the bath (electrical ground). Two voltage protocols were used to study the equilibration behavior of the system: Protocol 1 which applies consecutive voltage pulses either starting at positive voltage and incrementing towards negative voltage (starting voltage, V_s_, is greater than the end voltage, V_e_) or starting at negative voltage and incrementing towards positive voltage (V_s_ < V_e_). Protocol 2 applies a voltage (V_s_) for a variable duration and then switches to V_e_ for 10 seconds to record transient current. When V_s_ is positive, the pore is depleted of ions and therefore a greater portion of the current, upon switching to V_e_, goes towards maintaining concentration polarization. By varying the time spent at V_s_, the timescale of draining the pore of ions can be investigated.

Protocol 1 was used to investigate whether negative capacitance (*i.e.,* the magnitude of the current response) is dependent on salt concentration. Using V_s_=-1000 mV and V_e_=1000 mV, the capacitive current for V= -1000 mV was plotted on a logarithmic current axis (**Fig. 2a**). Noting that the resistive component was subtracted leaving only the capacitive current, negative capacitive current increases in magnitude with the KCl concentration. At the two lowest salt concentrations (10 mM and 30 mM), the first few milliseconds after the voltage switch also show positive capacitance before the slower (longer timescale) negative capacitance is observed. Conversely, the high salt concentrations are characterized by purely negative capacitance. As mentioned previously, negative capacitance is characterized by a series of exponential decays with a slowly decreasing time constant (*i.e.,* observed on a logarithmic scale as a non-linear slope over time).

**Fig. 2.**
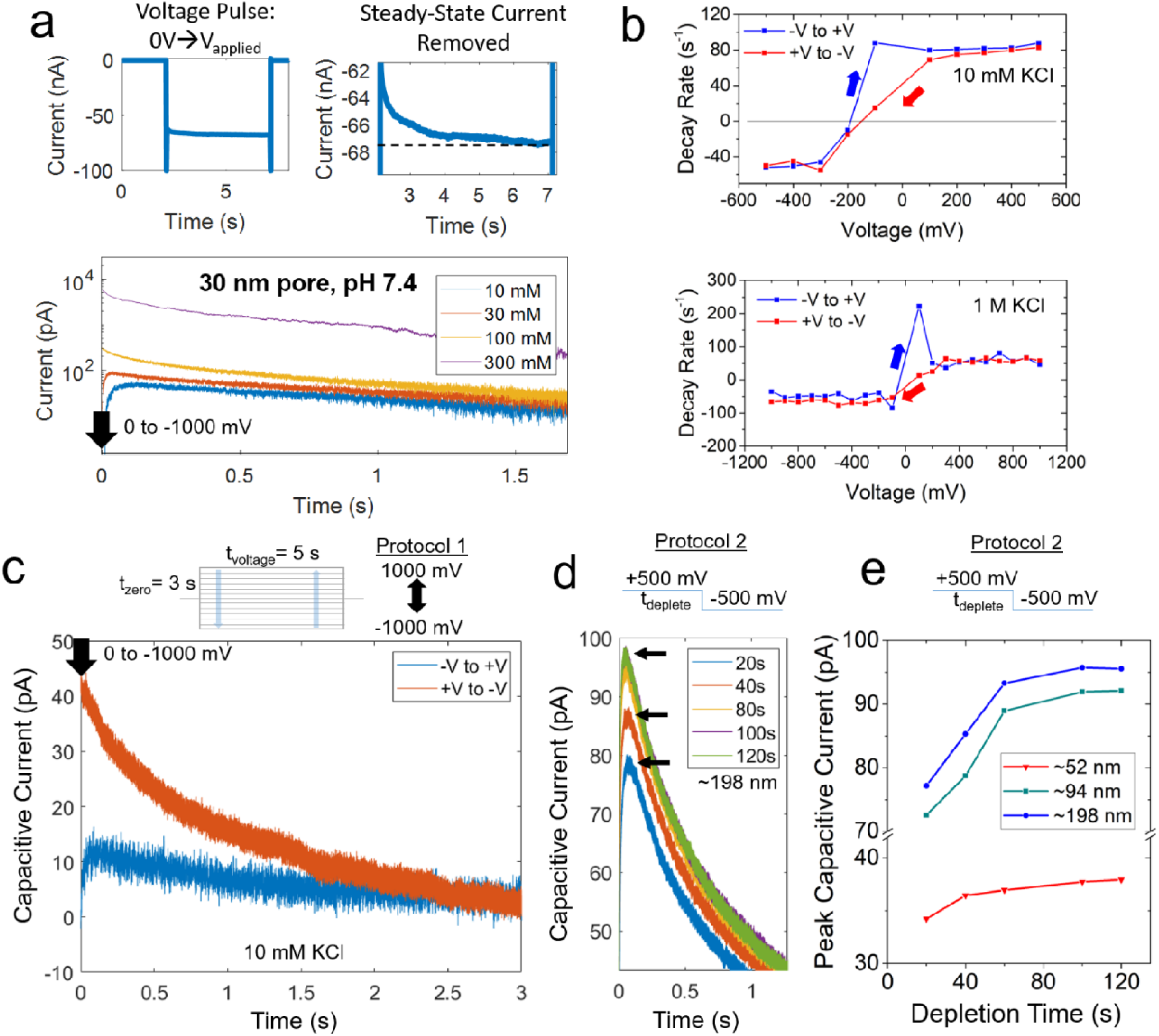
(a) Negative capacitive current response from a voltage pulse (0V to -1000 mV) recorded from the same pore (∼30 nm) at different salt concentrations: 10, 30, 100, and 300 mM KCl. (b) Extracted decay rates for negative capacitance at low (10 mM) and high (1M) salt conditions as a function of voltage. (c) The negative capacitive current spike for the 0 V to -1000 mV voltage switch when Protocol 1 was executed starting from -1000 mV versus starting at +1000 mV. Although the voltage pulse was the same, the history of the voltage pulses change the capacitive current response. (d) Protocol 2 data wherein the voltage was kept at +500 mV for various durations and then switched to -500 mV. (e) Peak capacitive current (*e.g.,* shown in (d) using black arrows) as a function of ion depletion time (duration spent at +500 mV prior to voltage switch) for different pore sizes. All recordings were repeated with a minimum of three devices, and representative traces are shown here.

The initial decay rate (inverse of the RC time constant) was further analyzed as a function of pore size and salt concentration. The extraction of the decay rate, briefly, entailed extracting the first 20 ms after the voltage switch and subtracting the current trace’s final mean value (i.e., the resistive current) to yield only the capacitive current. The resulting current traces were fitted to exponential decays (*I = I_o_e^-rt^*) and the decay rate, r, was extracted. Since the decay rate changes over time, a longer duration (>20 ms) led to poor fitting to a single exponential decay. The same voltage and fitting protocols were used for all instances where the decay rate is reported. Keeping in mind that a negative decay rate represents negative capacitance, we observed a transition from positive capacitance to negative capacitance at approximately -200 mV for 10 mM KCl and 0 mV for 1 M KCl (**Fig. 2b**). It is likely that the high salt condition leads to higher fluxes and thus more immediate charging of the tapered region of the nanopipette. Conversely, the 10 mM KCl condition has reduced ionic flux and thus cannot maintain the accumulation of positive charge inside the nanopipette at the voltage bias of -100 mV. In addition to the transition point between positive and negative capacitance, the 10 mM KCl condition also had two different decay rates at positive and negative voltages: 88 s^-1^ at V=500mV and -52 s^-1^ at V=-500mV. Meanwhile, the 1 M KCl experiments led to the nearly identical decay rates at V=500mV and V=-500mV (56 and 55 s^-1^, respectively). We speculate that the asymmetry in decay rates at low salt are linked to the rectification of the nanopipette (rectification ratio = I_-500_/I_+500_ = 9.4) which is weaker at 1M KCl (rectification ratio = 1.3, **see Supplemental Information**). The second noteworthy observation is the hysteresis observed in **Figure 2b**; specifically, the decay rates have different values at the transition point depending on the voltage step direction (-V to +V, or +V to -V). Hysteresis in the decay rates are attributed to mem-capacitance.

In addition to the hysteresis effect observed in the decay rates, the magnitude of the capacitive current was also found to have memory of previous voltage stimulation. In **Figure 2c**, two V = -1000 mV traces were plotted for a ∼30 nm pore filled with 10mM KCl (resistive component was removed from each trace). By only changing the order of the voltage pulses (starting at -1000 mV and incrementing up, versus starting at +1000 mV and incrementing down), the magnitude of the negative capacitive current also changed despite the pore, salt conditions, and voltage step were the same (*i.e.,* the only difference being the history of voltage pulses). Although only V=-1000 mV is shown here, all negative voltages changed their capacitive current magnitude depending on the history of stimulation (**see Supplemental Information**). These results were the clearest observation of “memory” within the nanopipette system and is a characteristic of the negative capacitance phenomenon.

Given that applying a positive voltage seems to enhance the negative capacitance effect, the time spent at positive voltage was varied to establish the nanopore mem-capacitance effect. Using Protocol 2, the voltage was kept at +500 mV for a certain duration (t_deplete_), followed by switching the voltage to V = -500 mV **(Fig. 2d**). Since the change in voltage is the same, the capacitive current should be the same if there are no memory effects. Instead, applying positive voltage prior to the voltage switch significantly affects the magnitude of the capacitive current but not the decay rate. The peak capacitive current (shown in **Fig. 2d** with black arrows) reached a maximum value after a depletion time of 100 s indicating that the ionic state of the pore has “memory” for over a minute. The nanopipette size dependence was also investigated by using the following approximate pore sizes: 52, 94, 198 nm (**Fig. 2e**). Once again. by varying t_deplete_, it was found that the nanopipette reached peak capacitance at 100 s across all the pore sizes tested (peak capacitance did not change between t_deplete_ = 100 s and t_deplete_ = 120 s). Although the pore size affects the magnitude of the capacitive current, memory of previous voltage stimulation has minimal pore size dependence (all pores had a memory effect of approximately 100 seconds).

Regarding the long timescale of the capacitive current, two contributions can be considered: the charge relaxation time, which is found by calculating the Debye time 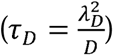, and the concentration diffusion time 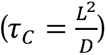^13^. For typical values of the Debye length (λ=1-10 nm) and diffusion coefficient (D=10^3^ µm^2^/s), the Debye time ranges from 1-100 ns. Conversely, we suspect that ions will need to diffuse several hundred microns in order to dissipate their charge^14^. As an example of the confinement experienced by an ion, consider a 10 nm pore with a typical outer cone angle of 10 degrees. Even if an ion travelled five microns inside the taper (starting from the tip), the inner glass confinement is still roughly 440 nm (*i.e.,* internal diameter). If a diffusional length, L, of 50 µm is used, the relaxation time would be 2.5 seconds which is roughly on the same scale as the current transients found here. This would also suggest that symmetric pores which do not have the tapered geometry would not display the same current transients during a change in voltage. Given the long equilibration times observed in the IV data, and the differences in the Debye time and the diffusion time, we conclude that the Warburg element is most applicable to nanopipettes and governs the long equilibration times observed within nanopipettes. Furthermore, if the ionic current is modelled as resistive (*i.e.,* steady-state) and capacitive (*i.e.,* transient), then the step-like change can be ignored and the capacitive current can be expressed as: 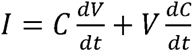. It is clear from this expression that either changes in the voltage or changes in the capacitance lead to transient current. Voltage-dependent changes in capacitance was first described in lipid membranes several decades ago^43^. We have observed here that these transient currents can persist for long periods of time (seconds) and cannot be ignored.

### Numerical Model

Studying non-linear electrokinetics using numerical models requires an important disclaimer about assumptions. The two critical assumptions are called the “constant field” and “electroneutrality” assumptions^44,45^. In fact, the electroneutrality assumption leads to paradox, or circular argument, in which the electric field is solved assuming no charge separation but the effect of the electric field induces charge separation which changes the electric field^46^. One of the main factors which govern violations in electroneutrality is the diffusion time which tends to be fast and quite capable of equilibrating unscreened charge^46^. However, as shown earlier, the diffusion time inside a nanopipette is slow and thus can lead to significant charge separation. With these caveats, we aimed to recapitulate our experimental results using finite element modelling keeping in mind that the most notable consequence of unscreened charge (charge density ≠ 0) is that voltage potentials within the ionic medium will develop and alter the voltage distribution within the nanopipette. For comparison, a neutral (i.e. zero surface charge) nanopipette was modelled and the voltage distribution along the axis of symmetry of the nanopipette shows no dependence on voltage polarity (**see Supplemental Information**); the spatial distribution of voltage was symmetric at both positive and negative voltage biases. As the surface charge was increased to -10 mC/m^2^, the voltage distributions become asymmetric with drastically different profiles at positive and negative biases (**Fig. 3a**). If the sensing zone is defined, albeit arbitrarily, as the region where 80% of the voltage drop occurs, positive voltages yield a smaller sensing zone while negative voltages yield a larger sensing zone. The distance over which the voltage drops can also elude to the strength of the electric field within the sensing zone; positive voltages having higher electric fields present inside the pore. At positive voltage, the voltage within the nanopipette is also shown to increase above the applied voltage (referred to here as the over-voltage condition^47^). At certain negative voltages (-200, -400, and -600 mV), the voltage also appears to go positive before returning to the negative voltage being applied. Although it is unclear whether the “over-voltage” conditions are occurring experimentally, the accumulation of ions/charge density is shown here to govern the voltage distribution and therefore the electric field inside and outside the nanopore. Therefore charge density, via Poisson’s equation, plays a significant role in sensing; especially since ionic current is measured by a voltage-sensing mechanism proposed earlier (**Fig. 1e**). Given the assumptions of electroneutrality, the quantitative accuracy of the results are still debatable, yet we believe they still provide insight into the qualitative behavior of the system and the origins of negative capacitance.

**Fig 3.**
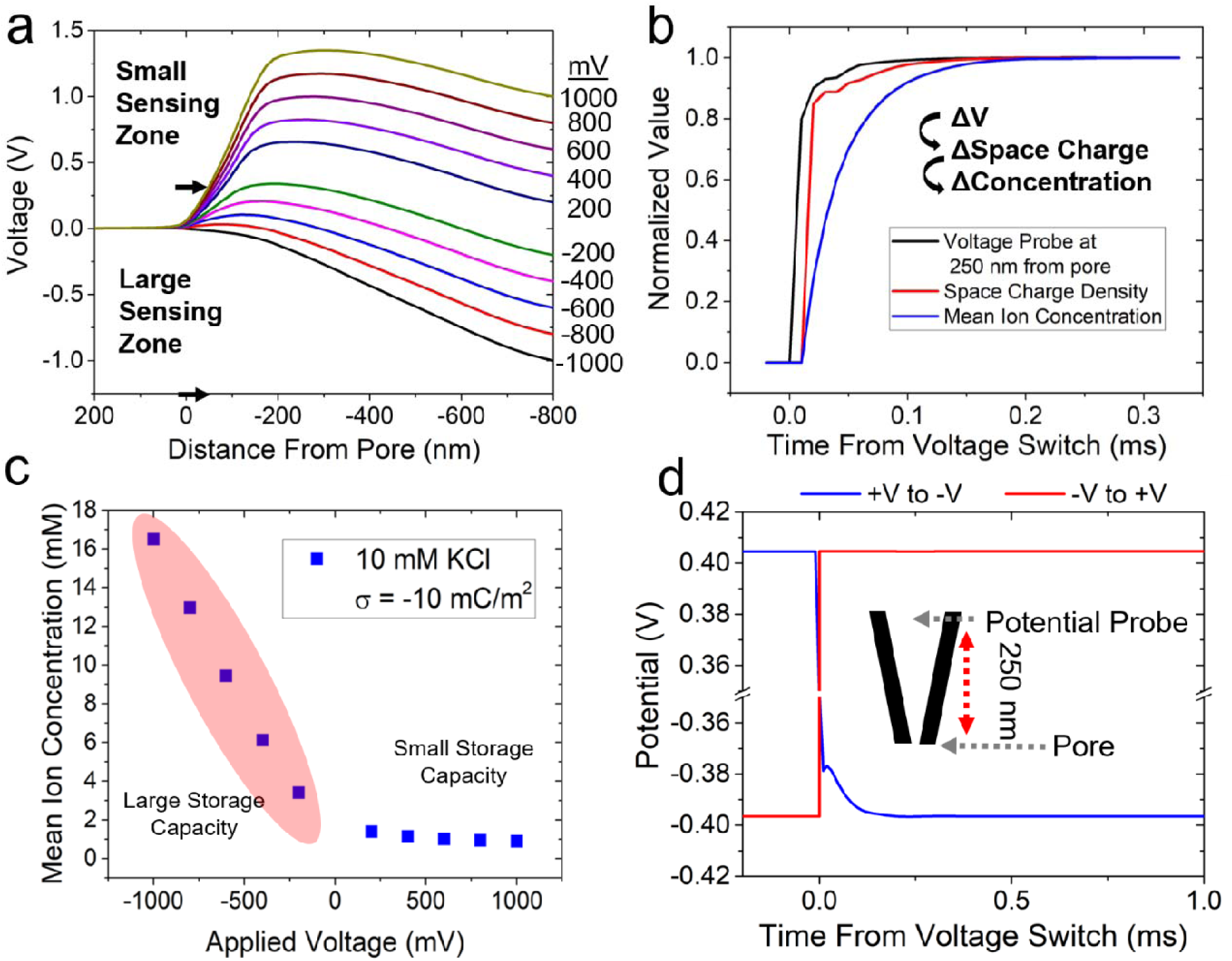
Finite element simulation results obtained by solving Poisson-Nernst-Planck and Navier-Stokes equations. (a) The spatial voltage distribution along the center (axis of symmetry) of the conical nanopore when the surface charge on the glass surface is -10 mC/m^2^. (b) Using the -10 mC/m^2^ condition, a time-dependent model was used to identify the causal relationship between voltage, space charge density, and concentration polarization. Each metric was extracted 250 nm from the pore (inside the taper). (b-c) The spatial distribution of voltage when the surface charge on the glass surface is increased to -5 and -10 mC/m^2^. (c) Mean ion concentration ([Cl^-^]+[K^+^])/2) as a function of voltage. The location of the concentration probe was 100 nm from the pore (inside the tapered region). (d) Potential probe plotted as a function of time from a voltage switch from +500 to -500 mV (+V to -V), and -500 to +500 mV (-V to +V). The potential probe was located 250 nm inside the taper.

Time dependent simulations are extremely useful to track changes in voltage within the nanopipette-ionic system since the deviation from assumptions can be observed in time and can provide insight into cause-effect relationships. Here, a time-dependent model (time resolution of 10 µs) was used to track the changes in voltage, space charge density, and mean ion concentration at a fixed position (250 nm into the nanopipette) during a voltage step from +500 mV to -500 mV. By normalizing the transient responses, it appears that the voltage change, as expected, precedes both any change to space charge density or ion concentration (**Fig. 3b**). Next, the space charge density is increased representing excess cations entering the taper. Lastly, the bulk concentration of ions increases as expected due to concentration polarization (noting negative voltages lead to higher electrolyte concentrations). Although concentration polarization has been well-documented, the driving force has not been proposed. Our results seem to suggest that voltage-driven excess charge (*i.e.,* non-zero charge density) is the driving force behind concentration polarization.

Non-linear electrokinetics is any process that depends nonlinearly on the applied voltage. Here, the mean ion concentration ([Cl^-^]+[K^+^])/2) inside the nanopipette is the first observation of non-linearity that is predicted by the finite-element model (**Fig. 3c**). The change in ionic concentration with voltage is critical to the observation of negative capacitance since voltage changes lead to a new steady state that needs to be reached. The simulation results show that the negative voltage regime is more dynamic in terms of its ability to store ionic charge. The ability to store charge differently at positive versus negative applied voltages implies that the capacitance of the Warburg element would be different depending on the voltage polarity. The polarity-specific effects of a voltage change was further characterized with a higher temporal resolution (1 µs). Using a voltage switch (V = +500 to V = -500 mV), a voltage probe located 250 nm into the nanopipette (centered along the axis of symmetry) showed that that there is an equilibration time associated with the re-distribution of voltage across the resistors (R_W_ and R_pore_). Switching from negative to positive voltage, however, does not lead to the same effect suggesting that the time required to enrich the pore with ions is slower than the time required to deplete the pore of ions (**Fig. 3d**). The ionic flux measured across the orifice of the pore indeed shows that switching from positive to negative voltage causes the pore to slowly achieve a higher conduction state which leads to negative capacitance (**see Supplemental Information**). The opposite scenario (-500 mV to +500 mV) shows that the flux decreases to towards a lower conduction state which mimics positive capacitance. The length of the transition period is also shown to be governed by the diffusion coefficients of ions within the tapered region (**see Supplemental Information**). Finally, using potential probes as a way to model ionic current fluctuations measured from nanopores may be more accurate given the voltage-sensing mechanism of patch-clamp amplifiers.

### Single Molecule Translocation Experiments at Low Salt

The term resistive pulse sensing implies that the resistance of the nanopore changes during molecular translocations and the capacitive current change is negligible or zero. However, the tapered region of the nanopipette has been shown to be best characterized by a variable resistor and a capacitor. Given that molecular occlusion can feasibly alter the ionic equilibrium of the pore charging and discharging of a capacitor element could be observable in single molecule translocation data. In all the experiments conducted here, DNA produced current enhancements (rather than current reductions) which may be linked to charge accumulation inside the tapered region of the nanopipette^48^. If the equilibrium is perturbed, the discharge of current should exhibit a decay rate governed by the RC time constant of the tapered region. Additionally, once DNA enters the pore, the DNA itself could store additional ions. If a decay of ionic current is observed in translocation events, the RC time constant during DNA entry and exit (event rise and fall time) could elucidate molecular properties such as charge or the length of the molecule.

The prototypical nanopore experiment involves placing λ-DNA in *one* side of the nanopore (pore diameter in this experiment was approximately 20 nm, see **Supplemental Information**) and applying a voltage bias. Here, DNA was placed in *both* sides of the nanopore so that the voltage polarity-specific properties of the translocation events can be identified and linked to voltage-polarity specific negative capacitance (**Fig. 4a-b**). Interestingly, several notable observations can be made about the voltage-dependent data: (1) negative capacitance is observed at -500 mV and -400 mV, (2) DNA events are resistive at positive voltages and conductive at negative voltages, and (3) the DNA event signatures at -500 mV and -400 mV seem to be lowpass filtered beyond what is expected from the instrument lowpass filter. The first observation was reproducible and characterized all pores tested under 10 mM KCl. The second observation suggests that the event type (*i.e.,* resistive versus conductive) is either voltage dependent or translocation direction dependent. At positive voltages, the DNA outside the nanopipette translocates into the nanopipette. At negative voltages, the DNA inside the nanopipette translocates out of the nanopipette tip. Although the salt and DNA are the same, the event type is different for “in” versus “out” events. The prevailing theory that counterions on the DNA are responsible for conductive events seems unlikely since the DNA has the same counterions irrespective of the transport direction. Previous work proposed a pore-centric theory for the nature of conductive events which is based on a flux imbalance between cations and anions^48^; namely that excess cations going into the pore yields conductive events which indeed would occur at negative voltages. Furthermore, the concentration polarization effects described previously yield very different ionic concentrations and could explain the polarity-specific event types. The last observation (i.e., evidence of a time constant within each event signature) has not been described in the literature to-date but seems to coincide with the negative capacitance effect observed during a voltage switch (both occur starting at -400 mV). A time constant associated translocation events implies that there could be an ionic lowpass filter in addition to the lowpass instrument filter which is routinely used to reduce noise (**Fig. 4c**). Indeed, it was also clear from the current signatures (**Fig. 4d**) that if the event duration (*i.e.,* dwell time) was too short, the current enhancement would not reach its maximum value (*i.e.,* the events were attenuated in amplitude). Further information about the DNA event properties such as current modulation depth and dwell time can be found in the **Supplemental Information**.

**Fig. 4.**
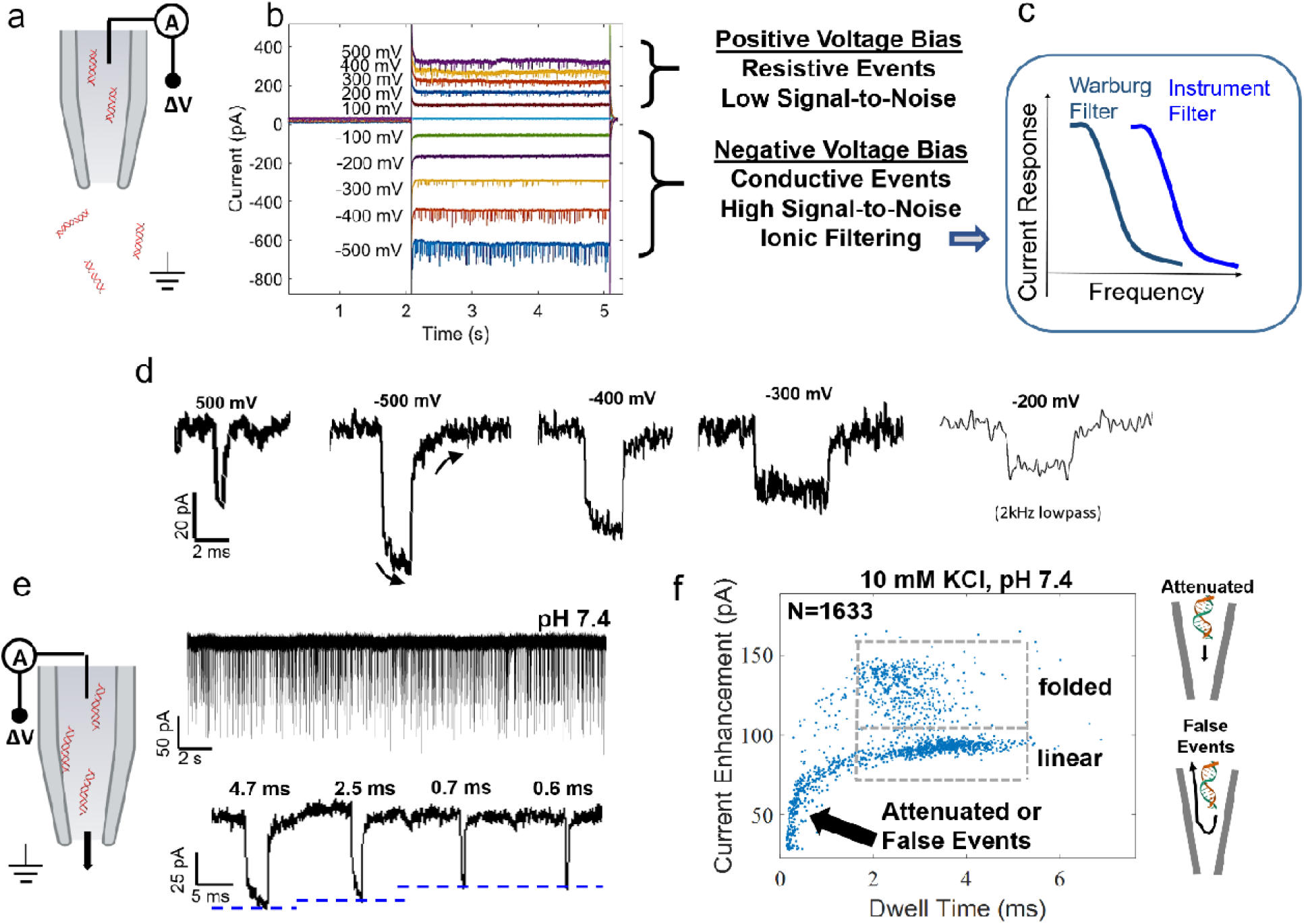
(a-b) Schematic of the experimental conditions (*i.e.,* DNA location) for data shown in (b). λ-DNA translocation data at a different voltages (+500 mV to -500 mV, in increments of 100 mV) for the following condition: 500 pM λ-DNA, 10 mM KCl (pH 7). (c) A schematic which illustrates the concept of Warburg filtering; specifically, that the ionic filter frequency is lower than the instrument lowpass filter. (d) Event signatures for the following voltages: +500 mV, -500 mV, -400 mV, -300 mV, and -200 mV. Unless otherwise noted, the instrument filter was a 10 kHz lowpass filter. (e) Current trace and representative DNA events. The analyte, λ-DNA, was only located inside the nanopipette with 10 mM KCl (pH 7.4) on both sides of th nanopipette and events were observed at negative voltages. The data in (e) was recorded at -400 mV. Event signatures, each labelled with their dwell time, to illustrate the attenuation of the current enhancements. (f) Scatter plot of event properties (peak current enhancement and dwell time) for λ-DNA inside the pipette and with 10 mM KCl (pH 7.4) on both sides of the nanopipette.

Since Warburg filtering and negative capacitance are only observed at negative voltage biases, DNA was placed inside the nanopipette for the rest of our experiments. At the pH of 7.4, events are slowed down by electroosmotic flow which opposes the electrophoretic transport of DNA. At this pH, event durations span a wider range compared to lower pH (for example pH 6, **see Supplemental Information**). It is evident that the dwell time of the DNA inside the pore has a direct impact on the current enhancement (**Fig. 4e-f**): the dominant event population which are labelled “linear” yield smaller current enhancements if the events are shorter in duration. At this stage, it is unclear whether events with smaller current enhancements are indeed attenuated or represent “false events”. False events are current modulations which do not ultimately result in DNA transiting the pore^49^. A voltage sweep (-200 mV to -900 mV) was performed to help elucidate which mechanism can best explain the data: attenuated events or false events (**see Supplemental Information**). Attenuated events should increase with voltage since the translocation time decreases with larger electrophoretic force whereas false events should decrease with increasing voltage since the ability to reverse from the pore is even less likely given the higher electrophoretic force. Our results indicate that the number of attenuated events increases with voltage which we suspect is due to a higher number of events being less than the critical dwell time (∼1.5 ms).

The theoretical basis for an ionic filter has not been established and therefore we sought to find the decay rate associated with DNA entering the pore and DNA exiting the pore. Once again, DNA was placed inside the nanopipette with 10 mM KCl (pH 7.4) on both sides of the pore (**Fig. 5a**). Using data obtained at a voltage bias of -500 mV, the event rise (DNA entry into the pore) and event fall (DNA exit from the pore) were fitted to an exponential. An average current trace across all events (n=625) representing the event rise and event fall is shown in **Figure 5b**. The event rise (DNA entering the pore) was decayed faster compared to the event fall (DNA exiting the pore); noting that the rise and fall designations are for conductive events. Less than 10% of events were filtered out due to variability in the traces which led to poor fitting of an exponential (e.g., **Fig. 5b** inset). The peak value for the decay rates were 5300 s^-1^ and 2500 s^-1^ for DNA entry and exit, respectively. Since the prototypical lowpass filter can be modelled as an RC circuit, and the time constant is equal to the product of the resistance and capacitance, the equivalent capacitance of the pore with and without DNA can be calculated. The resistance value of the pore was obtained from the I-V curve (negative voltages only due to rectification) and was found to be 1.3 GΩ. The resulting capacitance values were 123 and 304 fF for DNA entering the pore and DNA exiting the pore, respectively. Although the change in capacitance is clearly modulated by the presence of DNA, the ionic and/or molecular basis for the change in capacitance is unclear. The decay rate being faster for DNA events, compared to a voltage switch from 0V to -1V, for example, is likely due to the degree of perturbation of the ionic species within the nanopipette (*i.e.,* the ionic flux change due to DNA is small compared to the flux change due to a voltage switch).

**Fig. 5.**
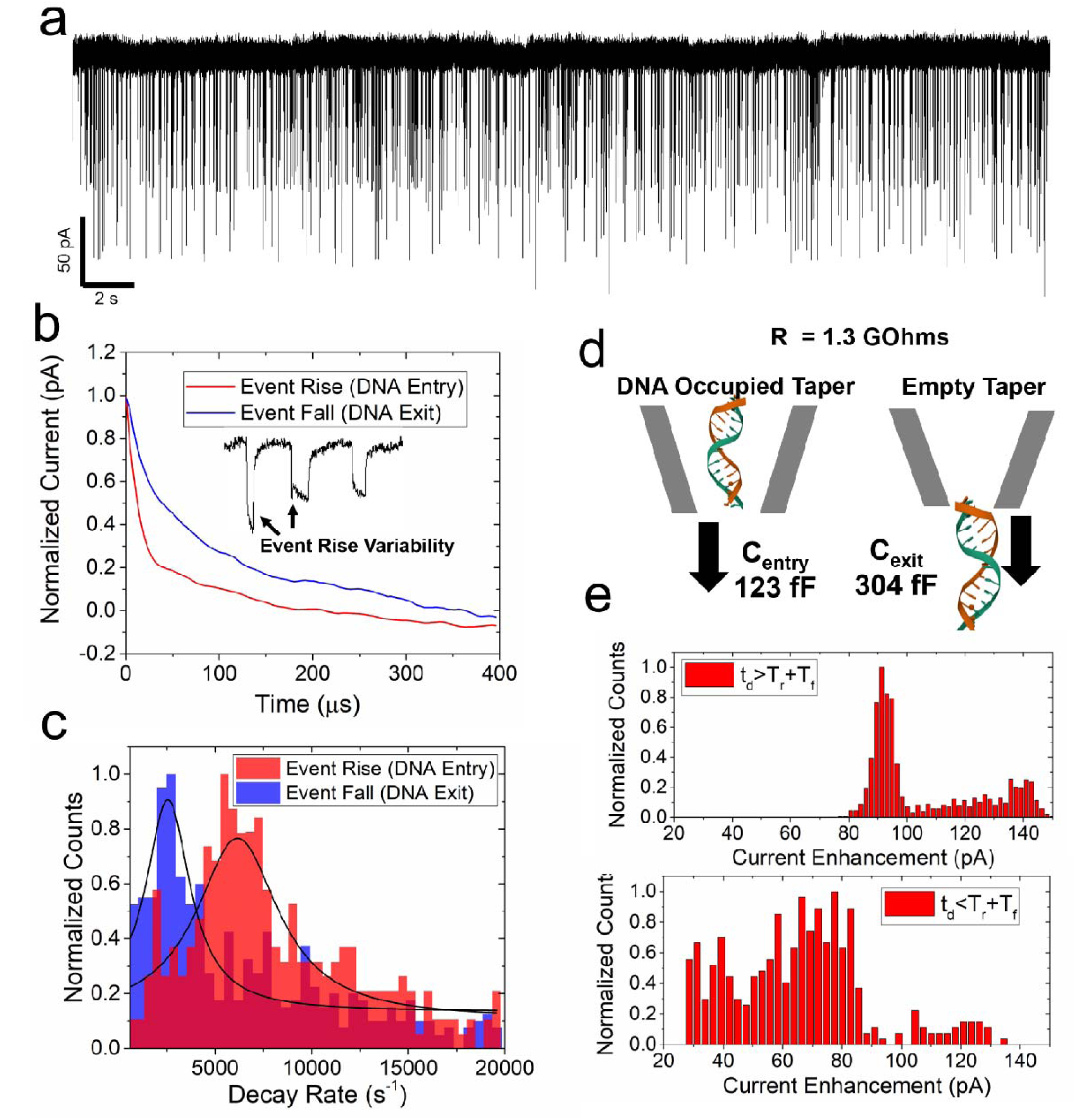
(a) A long (50 s) current trace showing conductive events (500 pM λ-DNA) with symmetric 10 mM KCl (pH 7.4). The applied voltage bias was -400 mV. (b) Average of different sub-regions of λ-DNA current signatures (event rise and event fall averaged over 625 events). Inset: examples of event rise variability wherein DNA folding or attenuation can hinder the observation of ionic current time constant. (c) Histogram of the time constant extracted from each event’s rise and fall. The ionic current decay towards its equilibrium value was fitted by an exponential and the time constant was extracted for each event (event rise data includes 625 events and event fall data includes 781 events). See Methods for more details. (d) Schematic illustration showing how the position of DNA inside the nanopipette seems to impact the measured capacitance (*i.e.,* RC constant) during the event rise and event fall. (e) Current enhancement histograms for events which have dwell times (t_d_) above (top) and below (bottom) the critical event duration (T_critical_=T_r_+T_f_). Below T_critical_, attenuation of the current signature is expected.

Warburg filtering only occurs at negative voltage biases and thus, assuming an analyte (such as DNA) is negatively charged, the analysis shown here is critical to when the analyte is placed inside the nanopipette. The polymer-electrolyte method of nanopore sensing, for example, requires the analyte to be inside the nanopipette^50,51^. The polymer-electrolyte method has many advantages however they generate conductive events that have the same exponential tail shown here. Other methods, such as hydrogel-filled pores, use a “load” and “eject” methodology to get DNA inside the nanopipette and observe similar event shapes as well^52,53^. Given the growing body of work that use non-standard electrolyte conditions, we sought to develop a theoretical framework for obtaining the minimum event duration that would result in no attenuation of the events (even when instrument bandwidth is sufficient). The equation that describes the relationship between the decay rate, time constant, and the effective rise time of a lowpass filter is: 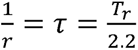, where T_r_ is the rise time (see **Supplemental Information** for derivation). Using the two values of the decay rate, the rise times of the effective filter were found to be 880 µs (event fall) and 415 µs (event rise). While a typical filter has the same time constant for both the rise and fall of an event (minimum event duration without attenuation is 2×T_r_), here the minimum event duration (T_critical_) without attenuation is the summation of the rise (T_r_) and fall (T_f_) times (880+415=1295 µs). Using this threshold value, events were split into two groups based on the value of the dwell time (t_d_). The current enhancements were distinctly different between the two groups (**Fig 5e**). The events with dwell times larger than 1295 µs showed two clear peaks corresponding to linear and folded DNA. The events that have dwell times less than 1295 µs has smaller current enhancements and no longer were able to resolve DNA folding. We have not yet investigated ways to eliminate Warburg filtering but speculate that it is intrinsically linked to the high signal-to-noise and conductive nature of events.

Attenuated DNA events are shown in **Figure 6a** and examples are shown for three categories of DNA events: linear, partially folded, and fully folded. A core assumption of the attenuation theory is that DNA events will be attenuated roughly proportional to the measured dwell time of the DNA events. The equation that can be used to correct for attenuation is:

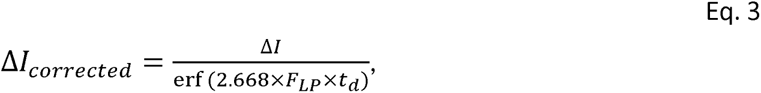

**Fig. 6.**
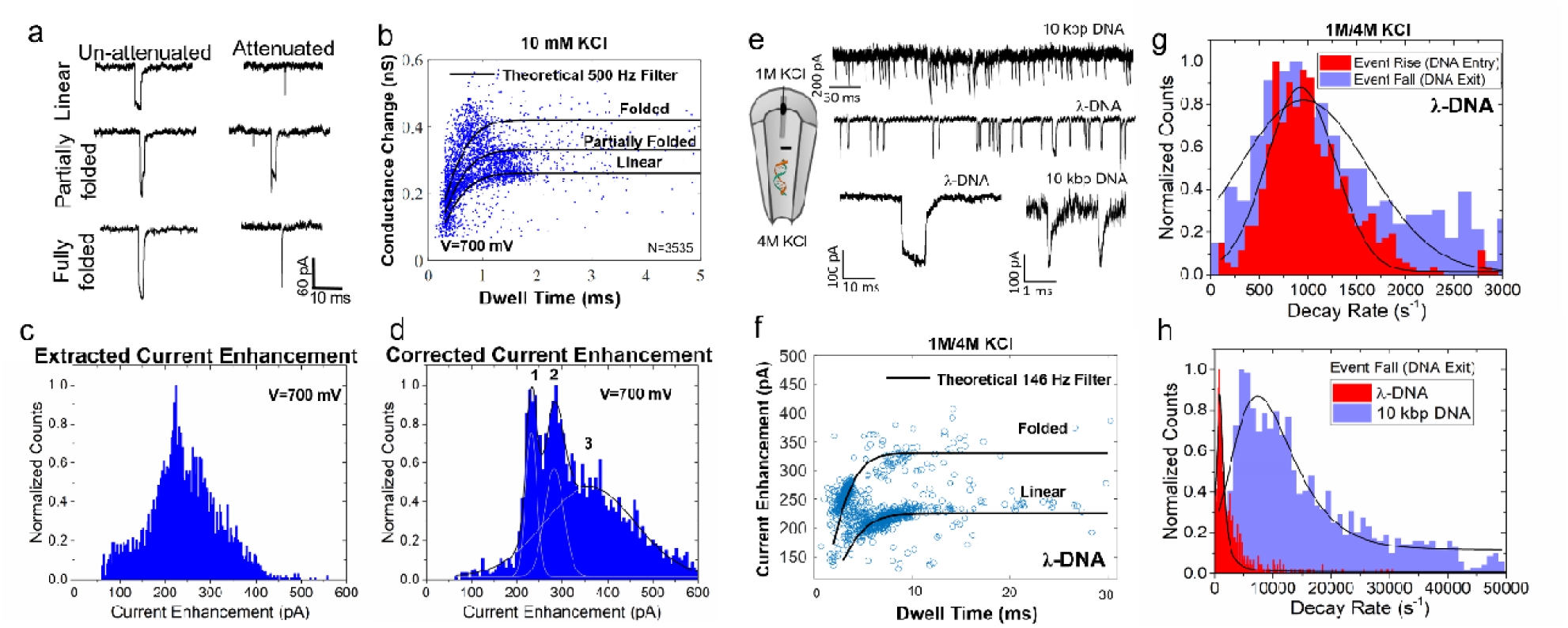
(a) Characteristic event signatures for linear, partially folded, and fully folded events (left: un-attenuated events, right: attenuated events). Events were recorded -700 mV (electrolyte: 10 mM KCl, pH 8). (b) The conductance change (ΔI/ΔV) for λ-DNA at voltage -700 mV (**see Supplemental Information** for other voltages). The black lines are the predicted attenuation curves for each DNA configuration assuming a cutoff frequency of 500 Hz. Note that all signals were recorded using a 10kHz lowpass filter. A total of 3535 events were recorded and analyzed in this dataset. (c-d) Extracted and corrected current enhancement histograms for the voltage bias of -700 mV. Gaussian peak fitting was performed on (d) and shows three peaks corresponding to linear (1), partially folded (2), and fully folded (3). (e) Current traces showing 10 kbp DNA and λ-DNA events collected under asymmetric salt conditions (1 M/4 M KCl, pH 7.4). All events were collected at a negative voltage bias (-400 mV) and events were conductive. (b) Decay rate histogram for both the event rise and fall which corresponds to the DNA entering the pore and exiting the pore, respectively. (c) Scatter plot of λ-DNA event properties and a theoretical lowpass filter with a cutoff frequency of 146 Hz. (d) Histogram of the event fall (DNA exit) decay rate for λ-DNA and 10 kbp-DNA. All data was collected at a voltage bias of - 400 mV.

where F_LP_ is the cutoff frequency of the lowpass filter. The error function alone can also be used to generate an expected attenuation curve for various dwell times. Experiments were performed for voltage biases in the range of -200 mV to -1000 mV (10 mM KCl, pH 7.4). The data for -700 mV is shown in **Figure 6b** however data for all voltages can be found in the **Supplemental Information**. The error function was then used to generate the expected attenuation curves for the three DNA states: linear, partially folded, and fully folded. For all three DNA configurations, the lowpass filter frequency (F_LP_) which best fit the data was 500 Hz. This effective lowpass filter frequency is significantly smaller than the 10 kHz instrumentation filter that was used to reduce electrical noise.

A histogram of the current enhancements was generated for both the extracted ΔI and the corrected ΔI based on Equation 1 (**Figure 6c-d**). The extracted ΔI histogram lacked distinguishable peaks corresponding to the different configurations of the DNA. Furthermore, the events below 150 pA are entirely made up of the attenuated events and do not fit will with a single Gaussian population centered at the first peak located at 179 pA. Using Equation 3, all ΔI values were converted to ΔI_corrected_. The correction yielded three easily distinguishable peaks corresponding to linear (1), partially folded (2), and fully folded (3) in **Figure 6d**. A cutoff filter frequency of 500 Hz was used for the correction.

### Molecule-Specific Decay Rates at Asymmetric Salt Conditions

At low salt conditions, the translocation of DNA yielded current enhancements which were characterized by two different decay rates towards their equilibrium current values (*i.e.,* the blocked current state, and the open pore current state). Another salt condition has also been found recently to produce current enhancements and occurs when there is a salt imbalance^54^. As opposed to 10 mM salt which exhibits strong electroosmotic flow that creates a flux imbalance, the asymmetric salt condition causes an asymmetric number of cations versus anions to enter the pore and can create non-linear effects^48^. Using 1 M KCl inside the pore (along with either 500 pM λ-DNA or 500 pM 10 kbp DNA) and 4 M KCl outside the pore, current enhancements were produced at negative voltages (*i.e.,* electrophoresis drives DNA out of the pore). Based on TEM images of nanopipettes pulled using the same protocol (**see Supplemental Information**), we estimate that the pore size for both experiments was approximately 20 nm. Translocation data was collected at a voltage bias of -400 mV for both DNA analytes and the current enhancements were 223 ± 35 pA and 235 ± 47 pA for λ-DNA and 10 kbp DNA, respectively (**Fig. 6e**). The dwell times at the same voltage condition were most different and had a more bimodal distribution. The most notable difference was that the peak corresponding to the fastest dwell times was located at 2.2 ms for λ-DNA and 251 µs 10 kbp DNA, respectively (**see Supplemental Information**). Given that λ-DNA is 48.5 kbp, it is not surprising that 10 kbp would have such a short dwell time within the pore.

Exponential fitting of each event signature’s rise and fall yielded nearly identical decay rates (924 s^-1^ for the event rise and 910 s^-1^ for the event fall) based on the peak of the Gaussian fits (**Fig. 6g**). The decay rates for both the event rise and event fall were nearly the same and was unique to asymmetric salt; this was not the case for 10 mM KCl where the decay rate was higher for the event rise compared to the event fall. Since the Debye length is larger at 10 mM KCl compared to 1 M KCl, we suspect that surface charge phenomena may be important to sensing the capacitance change due to a molecule. The second interesting observation is that the value of the decay rates is much smaller (*i.e.,* takes longer to decay) than 10 mM KCl. The lower decay rate suggests that there is also a lower lowpass cutoff frequency (F_LP_) associated with these recordings. With the RC time constant being the inverse of the decay rate, and using —, the T_r_ is 2.4 ms which yields a T_critical_ of 4.8 ms. Using the relationship and — solving for F_LP_, we obtain an effective lowpass cutoff frequency of 146 Hz (**Fig. 6f**). Plotting the theoretical attenuation curves along with the DNA event properties (scatter plot of dwell time and current enhancement), there is a general agreement between the experimental data and the bandwidth calculations.

The extraordinarily low F_LP_ values found here are far below the instrument bandwidth (10 kHz in all experiments). The Warburg filtering effect therefore may limit the detection capabilities of certain experiments; namely those producing conductive events. Smaller molecules like protein are most susceptible to bandwidth limitations since they tend to translocate much faster. The 10 kbp DNA translocations were analyzed and fitted using the same procedure as the λ-DNA. The decay rates were much faster compared to the λ-DNA with a peak decay rate of 7382 s^-1^ for 10kbp DNA (**Fig. 6h**). The higher decay rate means that smaller molecules have a higher effective bandwidth which is beneficial to sensing. Only the event fall could be fitted however since the initial moments of the event lacked a clear exponential rise. Using a decay rate of 7382 s^-1^, the calculated T_critical_ was found to be 596 µs. Since nearly all the events for 10 kbp DNA had a dwell time less than 596 µs, we suspect that most of the events were attenuated. The attenuation of events is likely the reason that the rising edge of the event could not be fitted to an exponential (*i.e.,* the event never reached the fully blocked current state).

## Conclusion

The IV data and translocation data provide strong evidence for the existence of a Warburg element in the equivalent circuit representing the nanopipette sensor. Given the slow equilibration time observed in the IV data, one-dimensional diffusion-limited transport of ions is most fitting to our ionic system. Two major repercussions have been described: (1) long transient currents observed during a voltage switch, and (2) Warburg filtering. Since the capacitive current can be expressed as 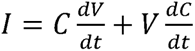, the mechanisms behind negative capacitance and DNA translocations can lead to a better mechanistic understanding ionic sensing. The results shown here are significant since the Warburg element can cause single molecule events, recorded by a nanopore, to be modulated and/or attenuated even when there is sufficient bandwidth of the ionic current amplifier. A purely resistive change due to molecular occupancy of the pore seems to be the least fitting model to explain the data shown here. Instead, there seems to be a resistive component as well as a capacitive component that appears only under certain experimental conditions. A Warburg impedance in series with the nanopore would provide a mechanism by which voltage fluctuations and/or capacitance fluctuations would vary depending on the degree of concentration polarization and accumulation of charge density. Although a Warburg element was modelled here as a single RC element, several RC elements would be needed to fully capture the variable decay rate of capacitive current. Therefore, multiple RC elements are needed to capture the exact nature of negative capacitance. The inclusion of non-standard electrical components could also be used to describe the current responses observed in this study. For example a variable resistor or a leaky capacitor would mimic and serve as a replacement for the RC element used here. The main issue with using a variable resistor is that the transition between states is not defined by the component itself. A leaky capacitor, however, is typically modelled as an RC element and therefore equivalent to a Warburg element with n=1 (n being the number of RC elements).

## Methods

### Nanopipette Fabrication

Quartz capillaries (Sutter Instruments, 1 mm outer diameter, 0.7 mm inner diameter) were pulled by a Sutter P2000 laser pipette puller. A relationship between conductance (using 10 mM salt) and pore size (via transmission electron microscopy: TEM) was used for nanopore sizing. See Supplemental Information for TEM images and corresponding conductance values.

### IV- and Gap-free Ionic Current Recordings

Current-Voltage (IV) relationships were recorded using an Axopatch 200B amplifier (Molecular Devices) and pClamp software. The IV protocol used episodic stimulation wherein the voltage sweeps started at zero voltage and initiated a voltage step at approximately 2 seconds. The duration of the file was adjusted to capture the dissipation of capacitive current. For IV curves that applied -1000 to +1000 mV, each sweep was extended to 10 seconds whereas IV curves which applied -500 to +500 mV, the sweep length was only 5.5 seconds. All IV curves started at the maximum negative voltage and made increments towards the positive voltage bias. All voltages are referenced to the outside of the nanopipette (*i.e.,* bath). The buffer used in all experiments, unless otherwise noted, was Tris:HCl with EDTA. For the IV curves conducted at pH 10, CAPS buffer was used instead of Tris. The λ-DNA and 10 kbp DNA used in experiments was obtained from New England BioLabs.

### Event Detection

The λ-DNA events were detected using custom MATLAB code which detects events based on a threshold value (the threshold was kept constant and was set as a multiple of the standard deviation of the background). The dwell time was measured as the full-width-half-max (FWHM) of the event signature. For example, if an event induced a current change of 100 pA (*i.e.,* peak current change), the time points where the event reached 50 pA current modulation were used to find the dwell time. All current enhancement values were the peak current enhancement of each event. No additional filtering of the events was done after event detection.

### Exponential Fitting and Time Constant Extraction

The time constants associated with a voltage switch as well as the DNA events were fitted by a single exponential of the form: Δ/ = Ce^-rt^, where C and r are fitting constants. If the exponential shape of the current transient was anything other than a positive current decaying towards zero, the signals were flipped (multiplied by -1) and the equilibrium current value was subtracted. All fits were checked for goodness of fit (R^2^>0.95). Unless otherwise noted, the first 20 ms of data (post voltage switch) were used for fitting. For fitting of DNA rise times (*i.e.,* the beginning of the event), the initial data point was the first FWHM point. The subsequent 0.8 ms of the event was then fitted to the exponential (Δ/ = Ce^-rt^) after subtracting the equilibrium value such that the current decayed to zero. Prior to this, only events >1 ms were used to ensure that 0.8 ms of data was obtainable for accurate fitting. For fitting the DNA fall times (*i.e.,* the end of the event), the second FWHM point was used as the initial data point. The subsequent 0.8 ms of data was then used for fitting. An illustration of the FWHM points used in the event rise and fall fittings are shown in the **Supplemental Information**.

### Finite Element Model

A full description of the finite element model as well as an exported report of the model can be found in the **Supplemental Information**.

## Acknowledgements

This work was supported by the Human Frontier Science Program (RGY0066/2018).

## Competing interests

The authors have no competing interests.

## Data availability

The data that support the findings of this study are available from the corresponding author upon request.

## Code availability

The code used to analyze data in this study are available from the corresponding author upon request.

## Notes

### Competing Interest Statement

The authors have declared no competing interest.

### Summary of Updates

New results are presented and described in the new manuscript.

