## Supporting Information for "Negative Mem-Capacitance and Warburg Ionic Filtering in Asymmetric Nanopores"

Table of Contents

I. Rectification in conical nanopores and negative capacitance at high salt (1 M KCl)
II. Finite element model
III. Pore sizing
IV. Negative capacitance at different voltages
V. Initial decay rates as a function of pH, salt concentration, voltage, and pore size
VI. Finite element model: neutral versus charged pore
VII. Finite element model: ionic flux timescale and diffusion coefficient dependence
VIII. Nanopore TEM images used in DNA experiments
IX. λ-DNA event properties in 10 mM KCl (DNA on both sides of pore)
X. λ-DNA event properties at lower pH (pH 6)
XI. λ-DNA event properties in 10 mM KCl (DNA inside the pore only): Voltage Sweep
XII. Derivation of the relationship between rise time and the RC time constant
XIII. Event properties for λ-DNA and 10 kbp DNA in asymmetric salt conditions
XIV. Event decays allude to current deficit and current excess
XV. Fitting exponential decays to events
XVI. Warburg variable impedance model

**I. Rectification in conical nanopores and negative capacitance at high salt (1 M KCl)**

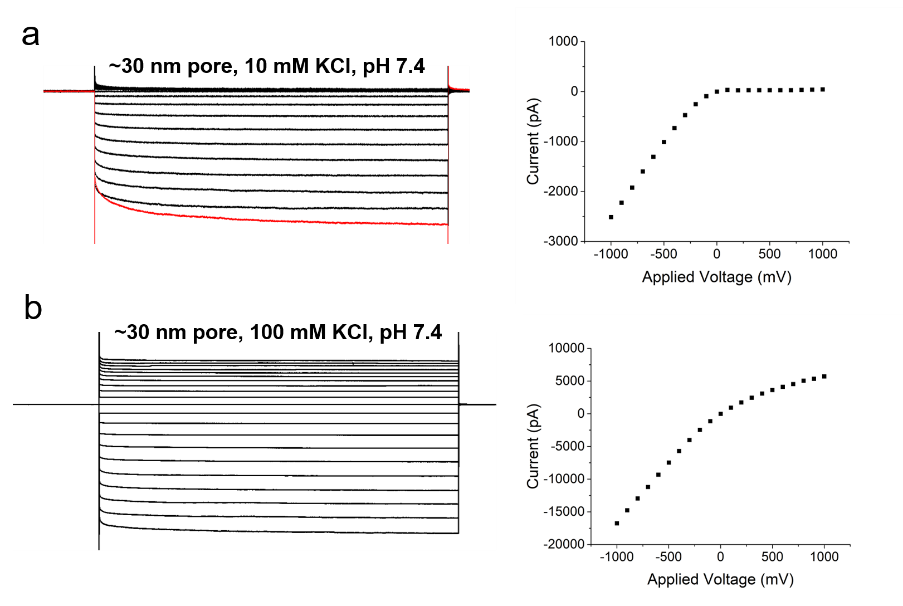

Supplemental Figure 1. (a-b) Current-Voltage (IV) response and curves for 10 mM and 100 mM KCl. The voltage sequence, unless otherwise noted, was always negative to positive (i.e. -1000 mV to +1000 mV) in increments of 100 mV. The same pipette pulling protocol was used in both experiments (pore diameter approximately 30 nm).

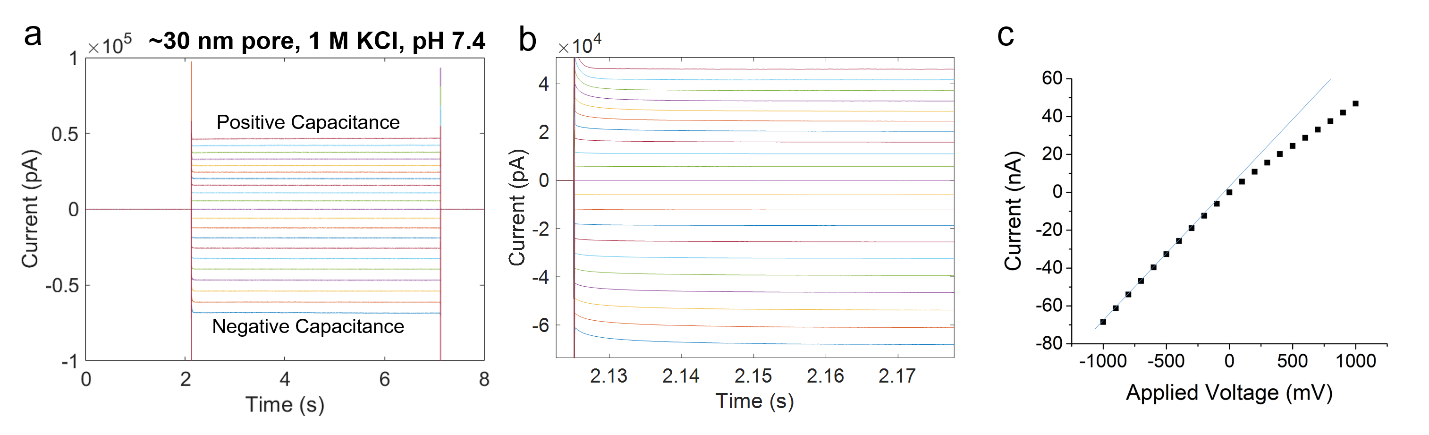

Supplemental Figure 2. (a) Full-scale version of the Current-Voltage (IV) response showing both positive capacitance (positive voltages) and negative capacitance (negative voltages). (b) Same data as in (a) but the time scale is shortened and zoomed into the first ~40 ms of the IV response. The decay of capacitive current is more clearly seen in (b) compared to (a). (c) IV curve extracted from the raw IV response data. The salt condition used in this experiment was 1M KCl, pH 7.4.

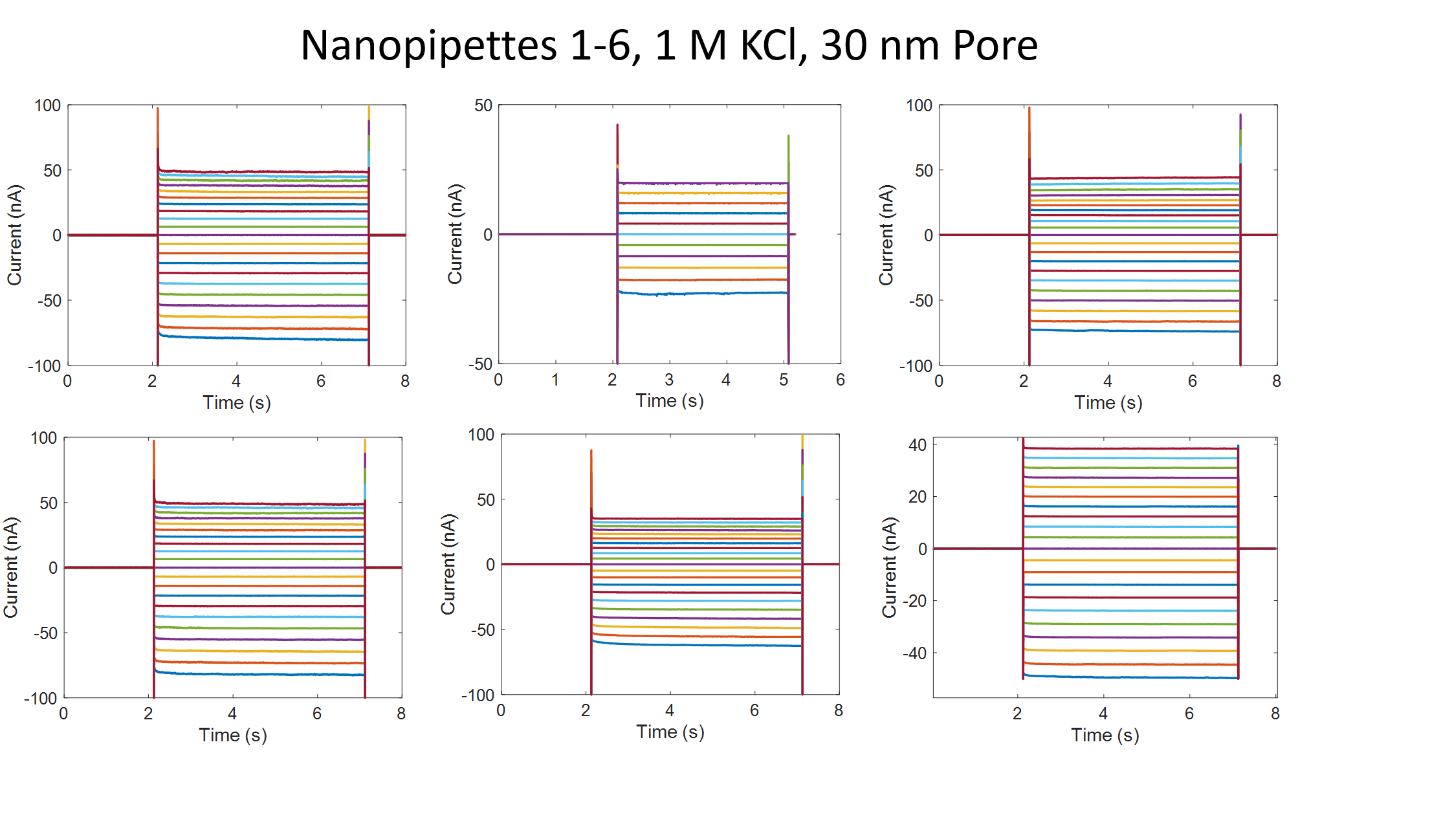

Supplemental Figure 3. (a) Repeats of the high salt condition (1 M KCl, pH 7.4) and a 30 nm pore. The IV curves were acquired from +1000 mV to -1000 mV (positive voltage applied first). The direction of the voltage pulses (+V to –V versus –V to +V) is essential to observe the negative capacitance phenomenon.

**II. Finite Element Simulations**

Finite element modelling was developed using COMSOL Multiphysics. The nanopores geometries were created based on the TEM images and pulling protocols achieved from the experimental studies. The simulation was designed at a low ionic strength electrolyte using the conical nanopore with a 25 nm diameter pore and a 4^o^ half cone angle. The diffusion coefficients were considered 2E-9 [m^2^/s] and 1.78E-9 [m^2^/s] for the potassium (K^+^) and chloride (Cl^-^) ions, respectively. The Poisson, Nernst-Planck, and Navier-Stokes equations were simultaneously solved to model the ionic behavior in a 2D axisymmetric steady-state model. The Poisson’s equation [∇^2^(V) = -ρν/ε] described the relationship between the electric potential and ion transport mechanism. The electrostatics boundary condition used for the glass was set at a surface charge density of -2E-2 [C/m^2^] in the vicinity of the pore opening to consider the surface charge contributions. The electric potential was set as variable field and the initial values were defined as zero potential. The space charge density was defined as ρ=N*e*z_1_*c_1_+N*e*z_2_*c_2_ for the electrolyte containing c_1_ (K^+^) and c_2_ (Cl^-^) ionic species where z and e were set as the valency and electron charge, respectively.

The Nernst- Planck equation was solved for the transport properties and ionic fluxes using convection, diffusion, and migration terms. The equation was described as: J_i_= -D_i_ ∇c_i –_ z_i_ μ_m,i_ Fc_i_∇^2^V where Ji, Di, c_i_, μ_m,i_ and z_i_ are the ion flux, ion diffusion coefficient, concentration, ion mobility and the charge number respectively. A no flux (J=0) condition was defined on the nanopore walls. The initial concentrations values of K^+^ and Cl^-^ species were set to 10E-3 [mol/L] for the entire domain. The electric force driven flow and pressure were modeled by the Navier-Stokes’s equation as: ρ(u. ∇)u = (-∇_p_ + η ∇^2^u -F (Σ z_i_ c_i_) ∇Φ. The u and Φ are the position dependent velocity field and potential field, z_i_ and c_i_ are species i charge and concentration in solution, ρ and η are the fluid density and dynamic viscosity, p is the pressure and F is the Faraday’s constant. Initial values of zero were assigned to the velocity field and pressure. The boundary condition for the pore wall was set to be u=0 (no-slip). To model the fluid flow, the volumetric force on the fluid was defined as ions space charge density multiplied by the electric field vectors. Pressure was specified as the boundary conditions for the inlet and outlet which in this work was set to zero.

*Note: A full COMSOL report of the simulations performed are also included as separate Supplemental Information documents.*

**III. Pore Sizing**

Using a transmission electron microscope, the size of each pore as a function of the “Pull” value on the P2000 pipette puller was tabulated and plotted. The low salt condition (10 mM) tends to complicate the sizing of nanopipettes due to the contributions of the debye layer and so a direct form of measurement like TEM was used rather than a theoretical equation. The conductance value of each pore was measured in 10 mM KCl (pH 7.4, Tris-HCl:EDTA) to form an empirical relationship between conductance and pore size.

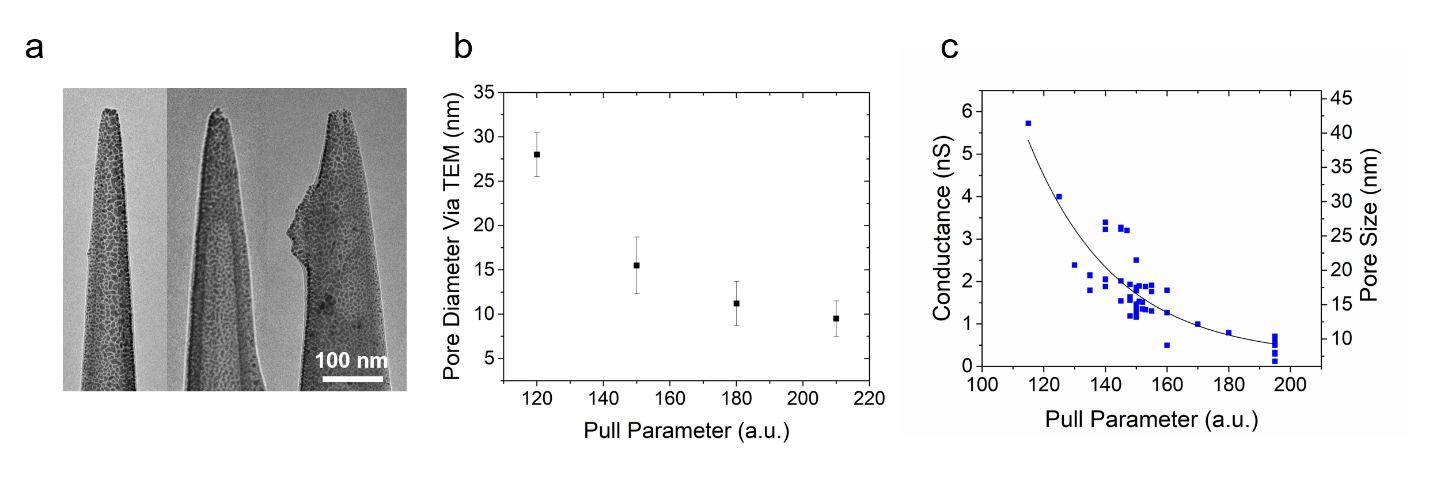

Supplemental Figure 4. (a) Transmission electron microscopy images for pipettes pulled at 210, 180, and 150 (left to right and in arbitrary units). (b) Pore size measured by transmission electron microscopy versus the pulling parameter on the P2000 pipette puller. The glass was Sutter Instruments 1mm O.D. and 0.7 I.D. quartz glass. (b) A plot showing the variability in conductance values acquired for each nanopipette pulled at a specific pull value. The salt used for conductance measurements was 10 mM KCl and Tris-EDTA buffer (pH 7.4).

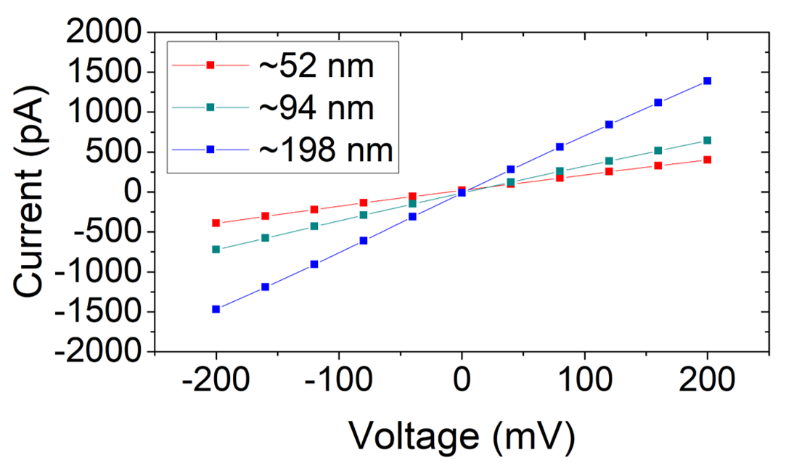

Supplemental Figure 5. Current-Voltage (IV) curves for three different pore sizes which were used to identify the pore size dependence of negative capacitance. The conductance of each pore was fitted using linear regression and pore size was calculated using the following equation:

$$d=\frac{4Gl}{\pi Kd_{b}}$$

where $l$ is the length of the conical pore (taper length), $K$ is the measured conductivity of the buffer, and $d_{b}$ is the diameter of the capillary (0.7 mm) at the beginning of the conical taper. The salt and buffer used in these sizing experiments was 1M KCl, Tris-EDTA (pH 7.4).

**IV. Negative capacitance at different voltages**

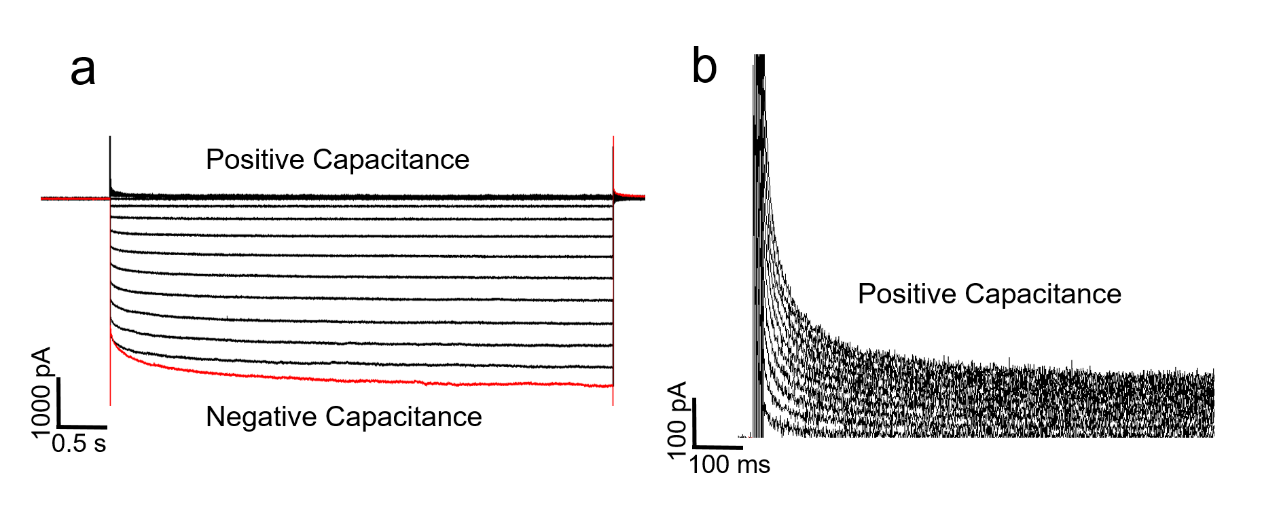

Supplemental Figure 6. (a) Current-Voltage (IV) response for a ~30 nm pore in 10 mM KCl, pH 7.4. (b) The positive voltages clearly show positive capacitance in the transient current response and rectification in the steady state current response.

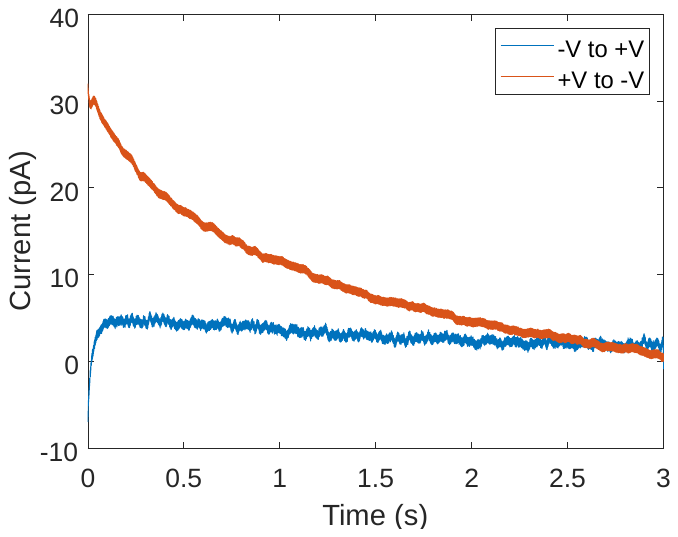

Supplemental Figure 7. (a) Comparison of negative capacitance transients when the Current-Voltage (IV) responses were conducted +V to –V versus –V to +V. The traces shown here are only the V= -700 mV responses with the resistive component subtracted, leaving only the capacitive transient. The time axis represents the time from when the voltage switch occurred (time zero is when the voltage was switched from 0 V to -700 mV). The pore was ~30 nm in 10 mM KCl, pH 7.4.

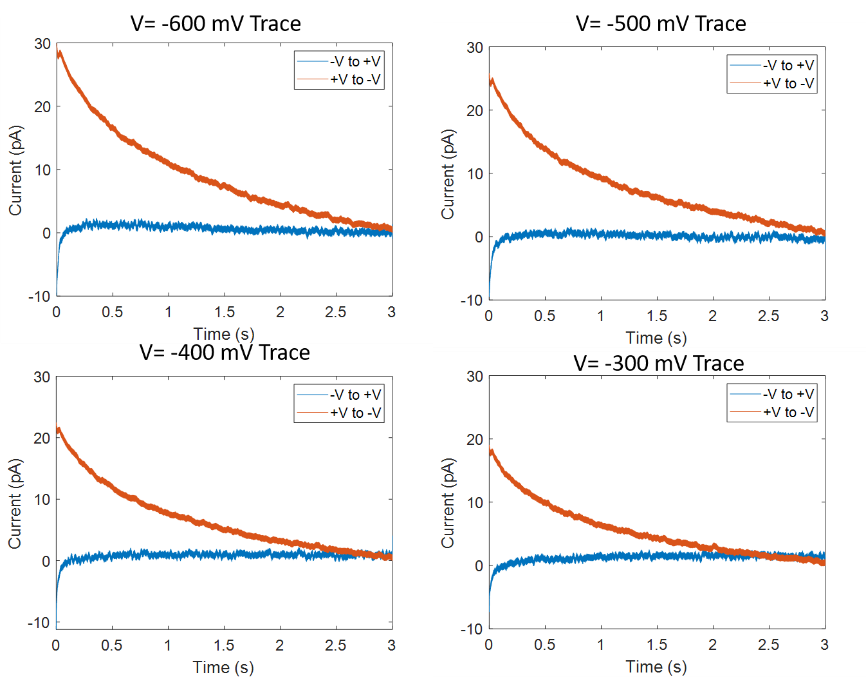

Supplemental Figure 8. (a) Comparison of negative capacitance transients when the Current-Voltage (IV) responses were conducted +V to –V versus –V to +V. The traces shown here are labeled at the top of each graph. The current responses have the resistive component subtracted, leaving only the capacitive transient. The time axis represents the time from when the voltage switch occurred. The pore was ~30 nm in 10 mM KCl, pH 7.4.

**V. Initial decay rates as a function of pH, salt concentration, voltage, and pore size**

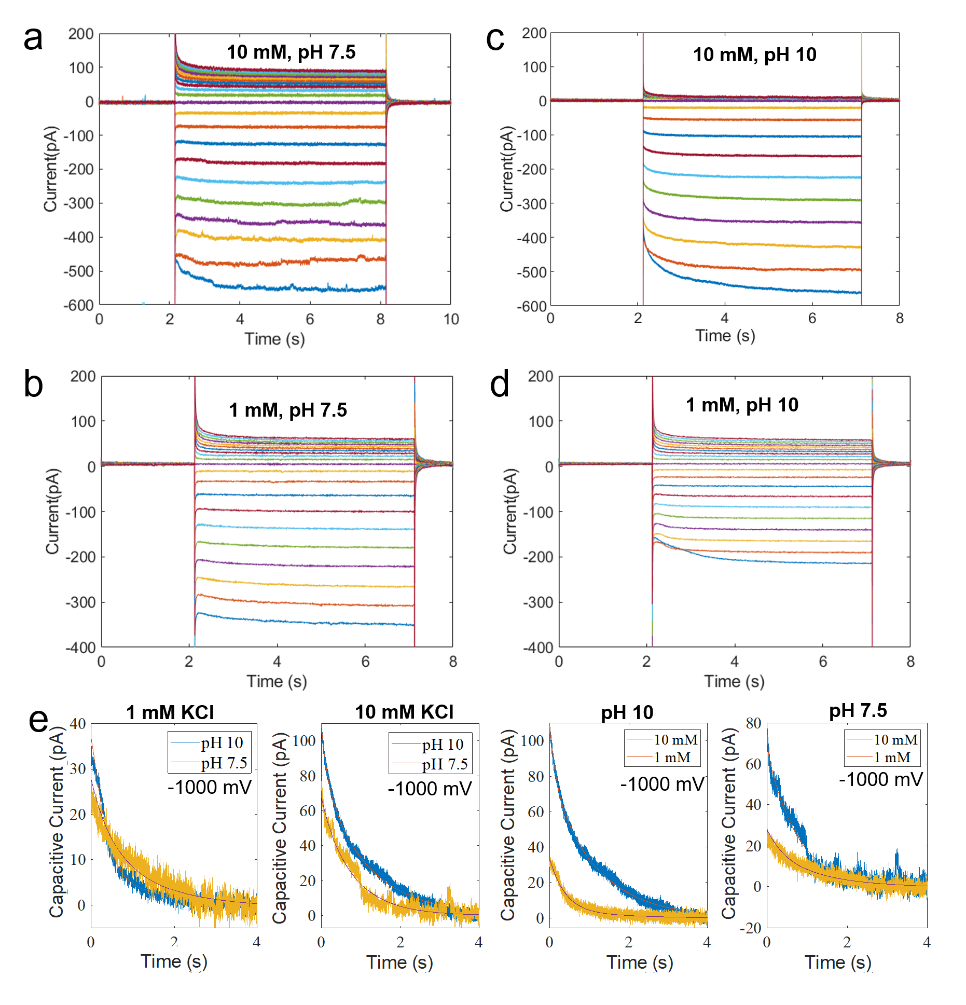

Supplemental Figure 9. Observations of negative capacitance as a function of salt concentration (1mM versus 10mM KCl) and pH (7.5 versus 10). All pores used were approximately 20 nm in diameter. (a-b) Current-Voltage (IV) responses at 10 mM and 1 mM KCl (pH 7.5). (c-d) Current-Voltage (IV) responses at 10 mM and 1 mM KCl (pH 10). (e) Capacitive current component only (resistive component was subtracted out) for all pairwise conditions. The voltage switch used for (e) was only the V=-1000mV voltage switch (time zero is when the pore bias was switched from 0V to -1000 mV).

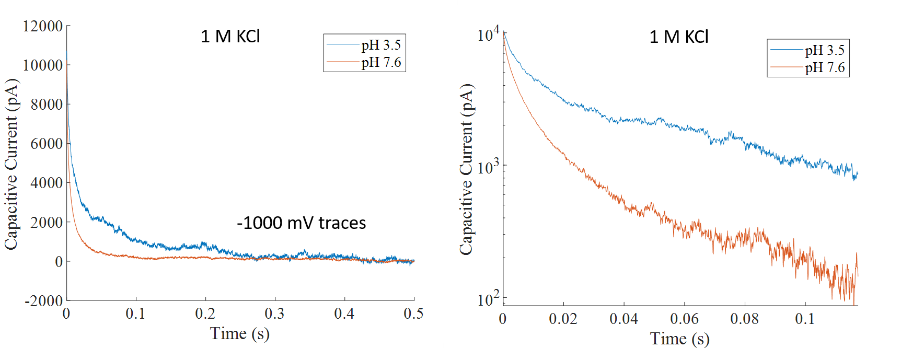

Supplemental Figure 10. Capacitive current responses (resistive component subtracted) for 1 M KCl (pH 3.5 and 7.6) and a pore size of approximately 20 nm. The voltage switch at time zero was from 0V to -1000 mV. The plot on the right has the y-scale on a log-axis.

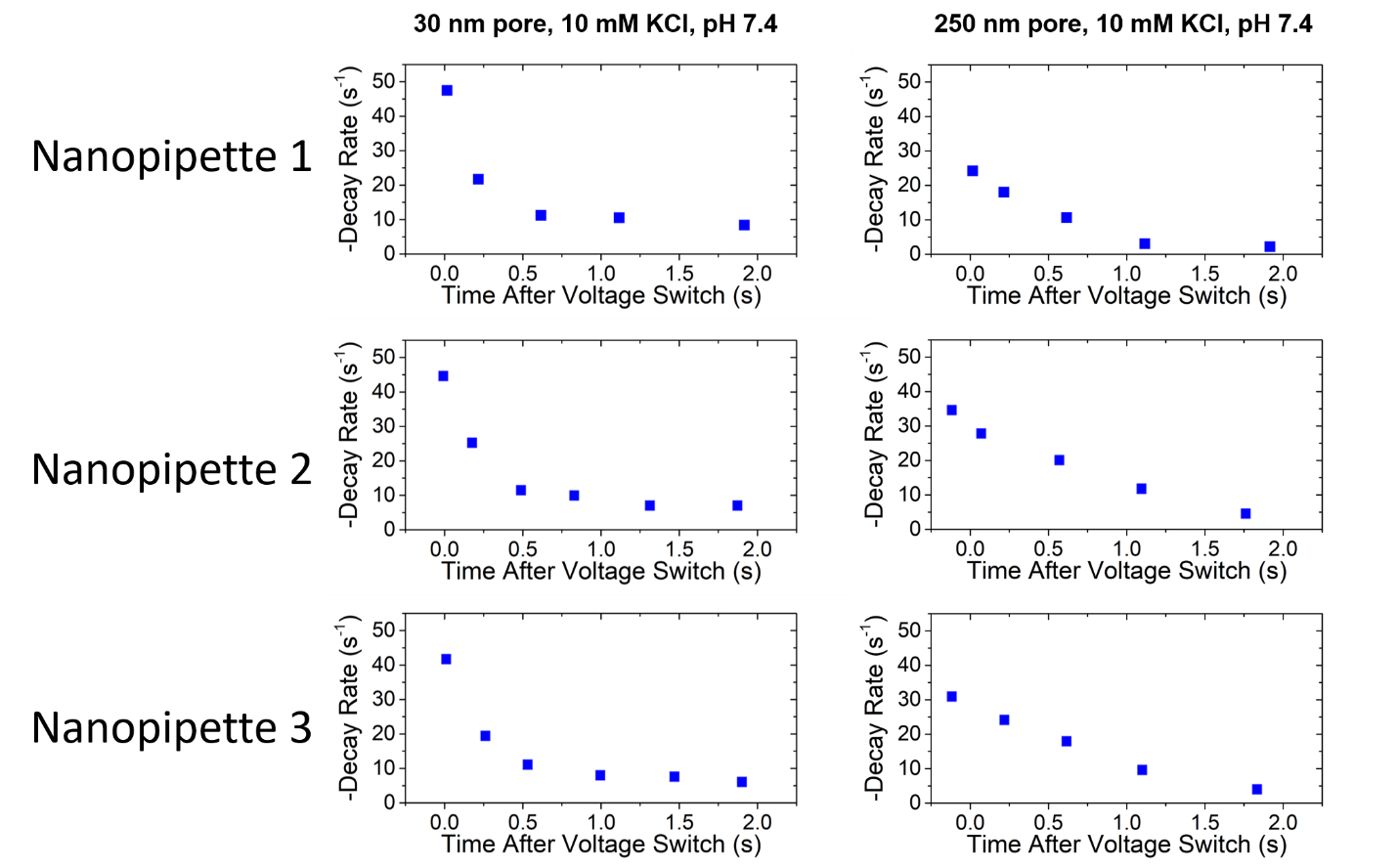

Supplemental Figure 11. Duplicate measurements across three nanopipettes from each category (30 nm and 250 nm; total of 6 nanopipettes). Each data point represents an the inverse of the RC time constant. The RC time constant therefore becomes larger and larger representing a more slowly changing current response. The decay rate was taken for a voltage pulse that starts at 0 volts and changed to V=-500 mV.

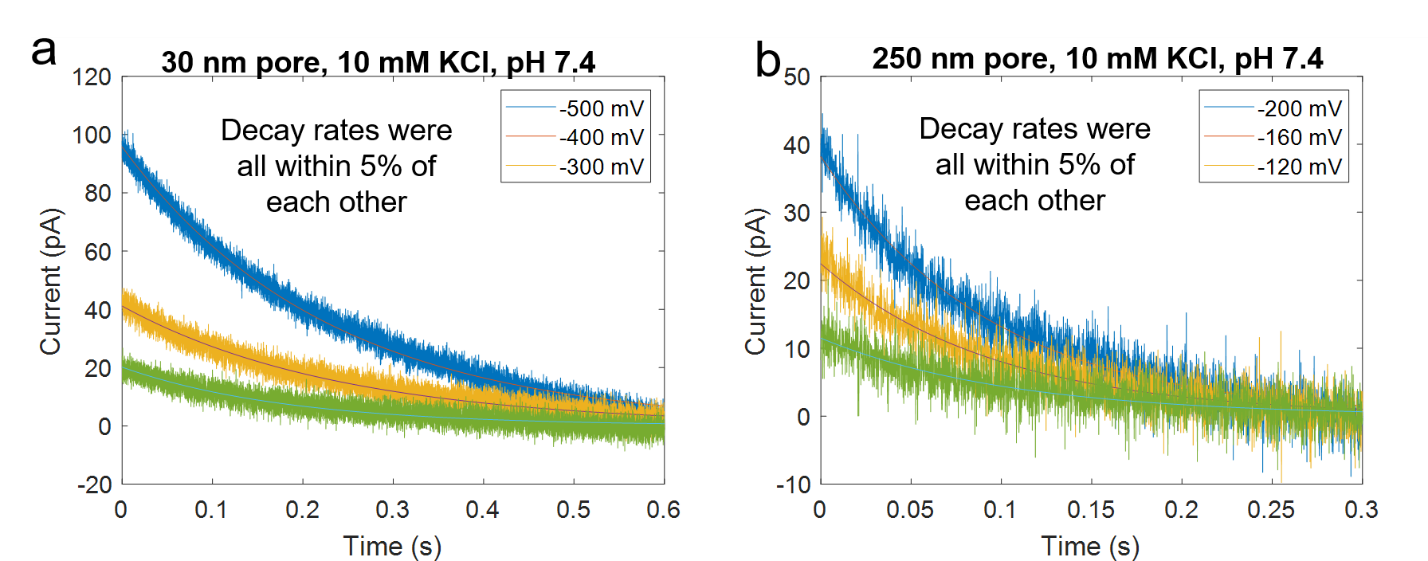

Supplemental Figure 12. Voltage dependence of the decay rate for a large (250 nm) and small (30 nm) pore. Although the magnitude of the capacitive current (i.e. transient) is voltage dependent, the decay rate was not affected. In each of the traces above, the same decay rate could be used to fit the data. If the decay rate was left as a free fitting parameter, the value would only vary (at maximum) +/- 5% from each other.

**VI. Finite element model: neutral versus charged pore**

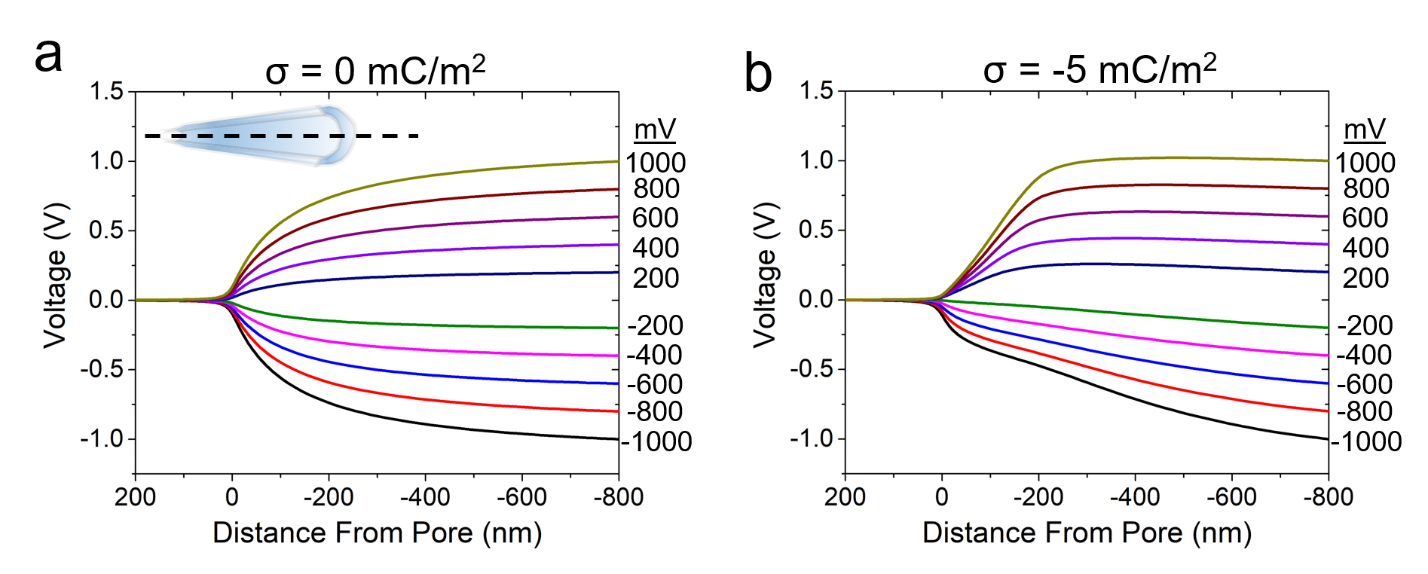

Supplemental Figure 13. (a) Finite element simulations for a pore (20 nm in diameter) that has zero surface charge. The voltage is plotted as a function of distance (in nm). With the bath side grounded, the voltage increases or decreases to the set point which is notated on the right side of the plot. The neutral pore shows symmetry at both polarities. (b) The voltage distribution inside a charged pore is asymmetric due to ionic charge dis-equilibrium.

**VII. Finite element model: ionic flux timescale and diffusion coefficient dependence**

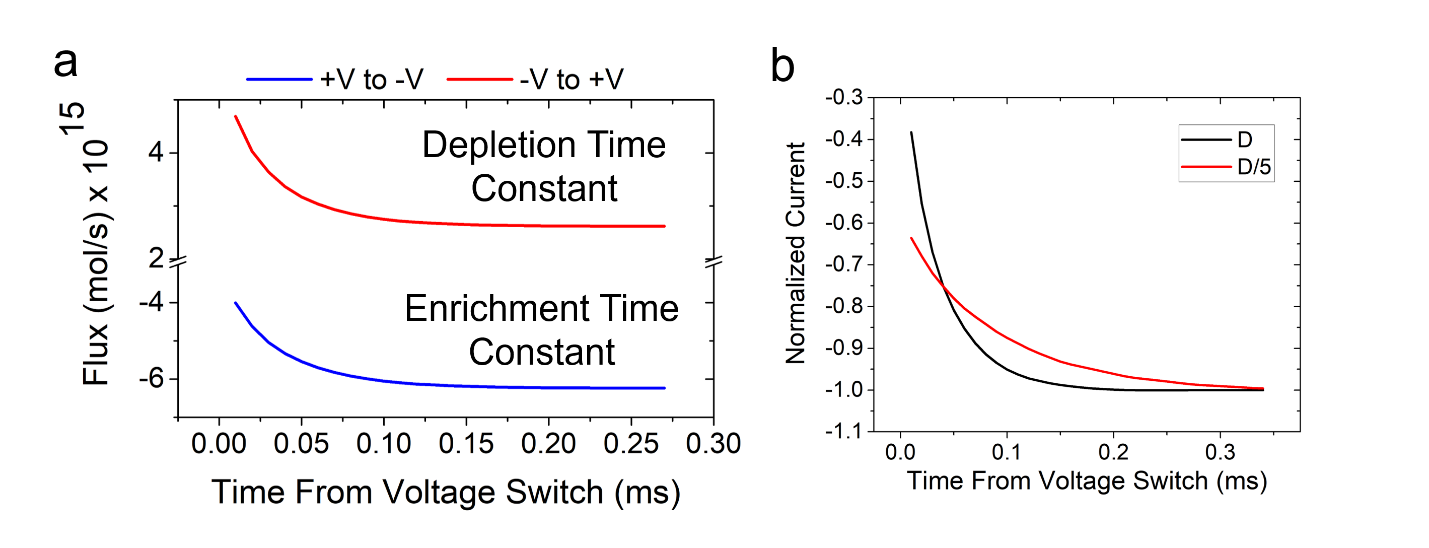

Supplemental Figure 14. (a) The voltage stimulus applied to the pore was +500 mV to -500 mV for the “+V to –V” condition and -500 mV to +500 mV for the “-V to +V” condition. For the “+V to –V” condition, the pore had to transition from a depleted ionic state (which occurs at +V) to an enriched state (which occurs at –V). It is noteworthy to show that the ionic flux is indeed smaller at t=0 compared to the equilibrium flux for the “+V to –V” condition. (b) Comparison of the normalized current response for the following conditions: (1) standard diffusion coefficients for K+ (2E-9 [m^2^/s]) and Cl- (1.78E-9 [m^2^/s]), and (2) smaller diffusion coefficients for K+ (0.4E-9 [m^2^/s]) and Cl- (0.356E-9 [m^2^/s]). The normalized current response (i.e. time constant) is critically dependent on the diffusion coefficients of the ionic species used in the simulation.

**VIII. Nanopore TEM images used in DNA experiments**

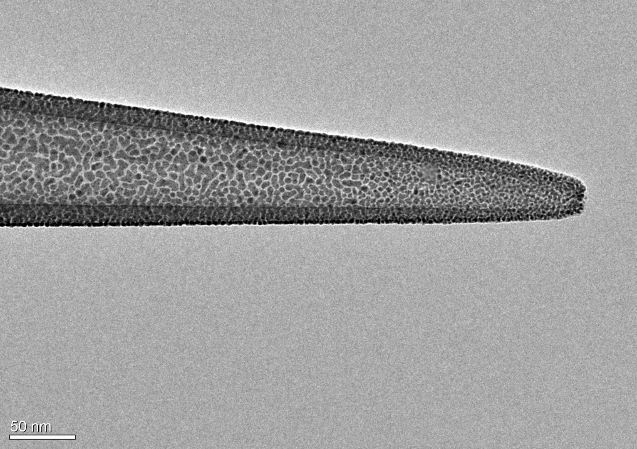

Supplemental Figure 15. Transmission electron microscope image for a quartz capillary (1mm outer diameter, 0.7 mm inner diameter) pulled to approximately 20 nm. The nanopipette was coated with 5 nm of gold prior to imaging to reduce charging and improve image quality.

**IX. λ-DNA event properties in 10 mM KCl (DNA on both sides of pore)**

The typical nanopore experiment involves placing λ-DNA in one side of the nanopore and applying a voltage bias. Here, DNA was placed in both sides of the nanopore and an IV curve was recorded. Not only is negative capacitance observed at the negative voltages, but the DNA produces a current increase as well (i.e. a conductive event). Even more surprising is that at positive voltages, DNA translocating the opposite direction yields resistive events. The event signatures themselves are uniquely different as well. Conductive DNA events appear to have a clear exponential decay/growth whenever the current changes. The existence of a leaky capacitor within the nanopore system could be useful to explaining these otherwise anomalous event characteristics. We further propose that charge accumulation and recombination of dissimilar ions can lead to current enhancements.

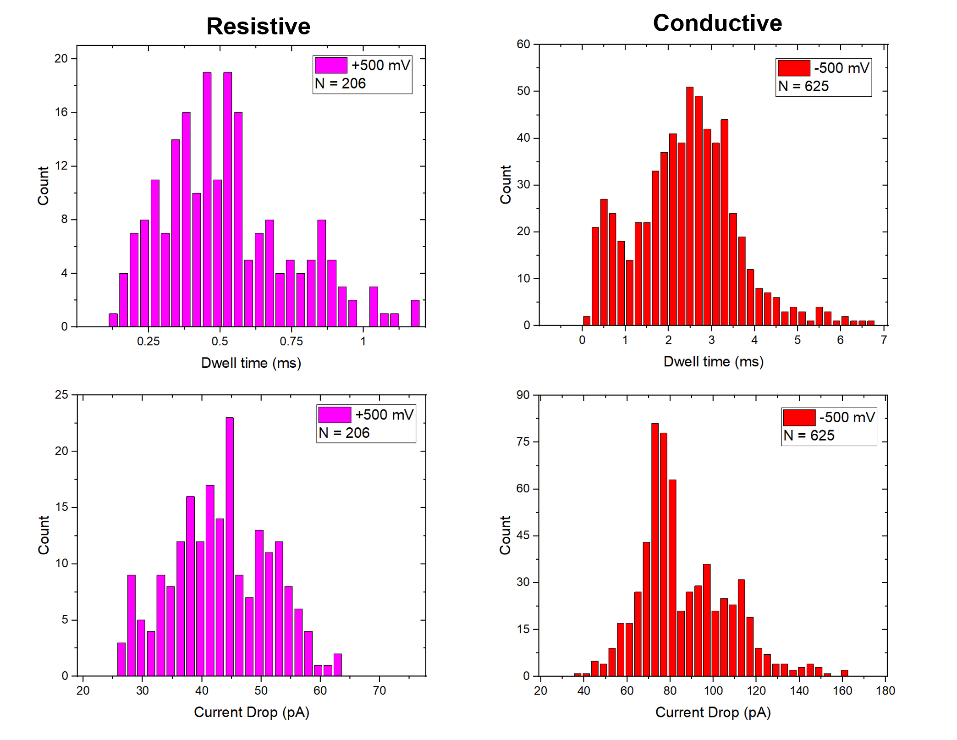

Supplemental Figure 16. DNA event properties (current modulation and dwell time) for λ-DNA with an applied voltage of -500 mV and +500 mV. At +500 mV, the events were resistive and has a reduced magnitude compared to their conductive counterparts (i.e. all events at -500 mV were conductive and had a higher amplitude). The resistive events also were faster (shorter dwell time) compared to the conductive events.

**X. λ-DNA event properties at lower pH (pH 6)**

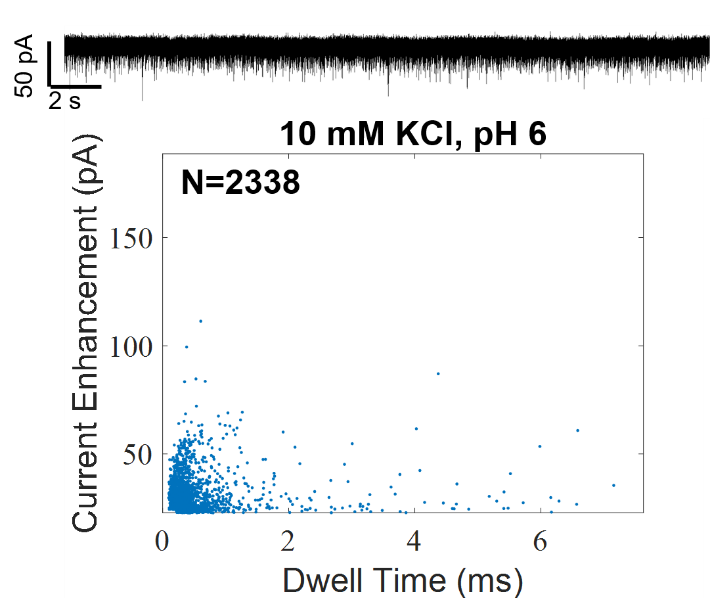

Supplemental Figure 17. λ-DNA translocation data under a voltage bias of -400 mV and in 10 mM KCl (no buffer). The pH was adjusted to pH 6 using dilute hydrochloric acid. The pore size of the nanopipette was approximately 20 nm and the DNA was placed inside the nanopipette so that the DNA would translocate out of the tip and into the bath. The dwell times are notably shorter than experiments at pH 7.4 which may be due to stronger electroosmotic flow which counters the electrophoretic force on the DNA. Due to the shorter dwell times, we seem to only collect attenuated events which do not reach their full amplitude. As a result, we also do not see any clear indications of DNA folding, which was clearly observed at pH 7.4.

**XI. λ-DNA event properties in 10 mM KCl (DNA inside the pore only)**

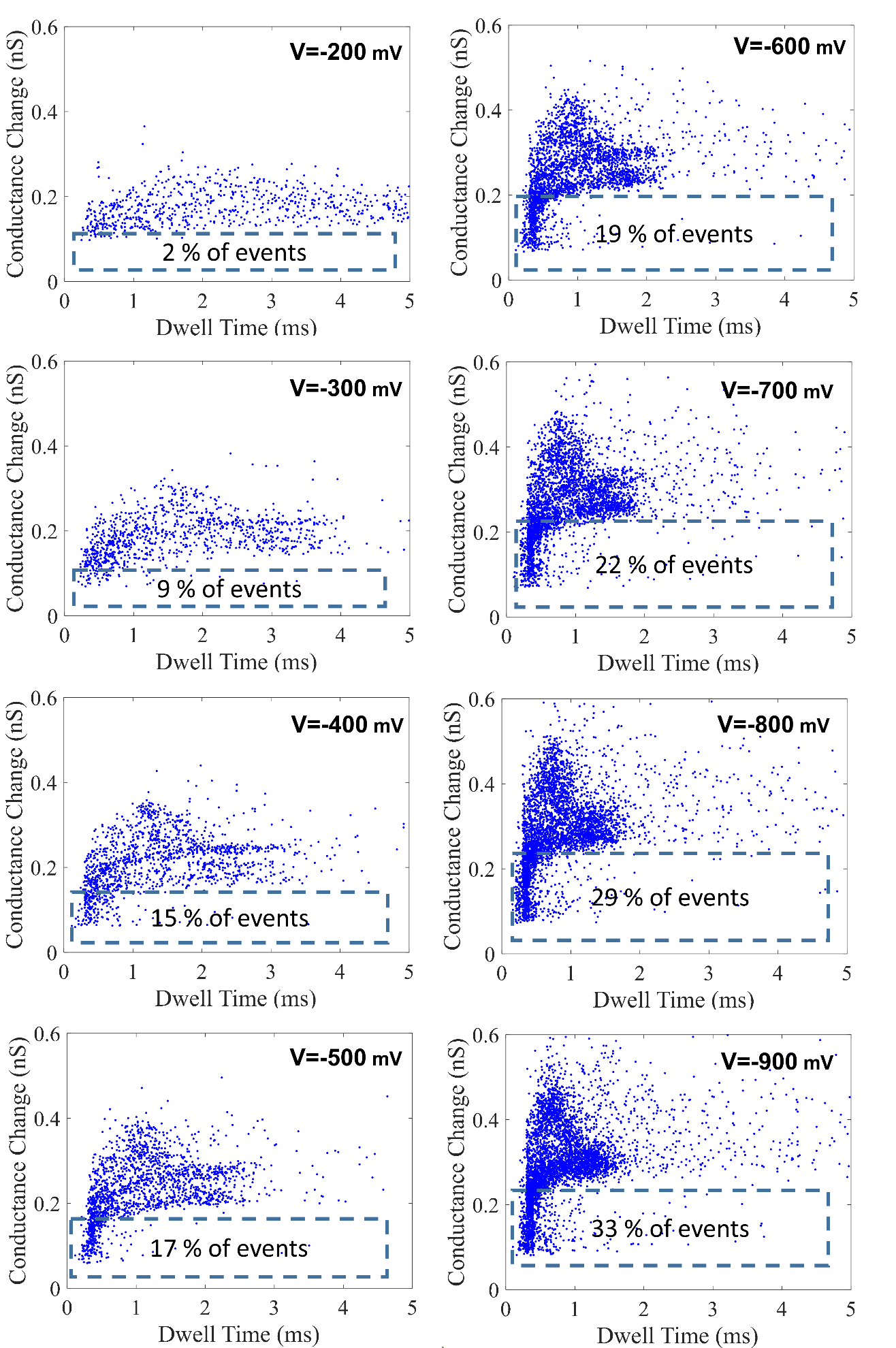

Supplemental Figure 18. Voltage dependent changes in event properties for λ-DNA in the low salt condition (10 mM KCl, pH 8). λ-DNA was placed inside the pipette and was translocated outwards. All voltages shown are negative relative to the grounded bath (external to the nanopipette). Current enhancements (in pA) were divided by the applied voltage to get the change in conductance (in nS).

**XII. Derivation of the relationship between rise time and the RC time constant**

For a step function that has an amplitude of 1 V, the time dynamics of the voltage across the capacitor is given by: $V(t)=1-exp(\frac{-t}{RC})$. The time constant is refined as RC and has units of time. The rise time of the RC lowpass filter is defined as the time it takes for the voltage to go from 10% (0.1 V) to 90% (0.9 V) of the final voltage (in this case 1V). Assuming the voltage switch happens at t=0, then the time to reach 0.1 and 0.9 is given by:

$$t_{0.1}=-\ln\left( 0.1-1 \right)*RC$$

$$t_{0.9}=-\ln\left( 0.9-1 \right)*RC$$

Since the rise time is equal to t_0.9_-t_0.1_, then the relationship between the rise time (T_r_) and the RC time constant is given by:

$$T_{r}=\left[ -\ln\left( 0.9-1 \right)+ln(0.1-1) \right]*RC=2.2*RC$$

**XIII. Event properties for λ-DNA and 10 kbp DNA in asymmetric salt conditions**

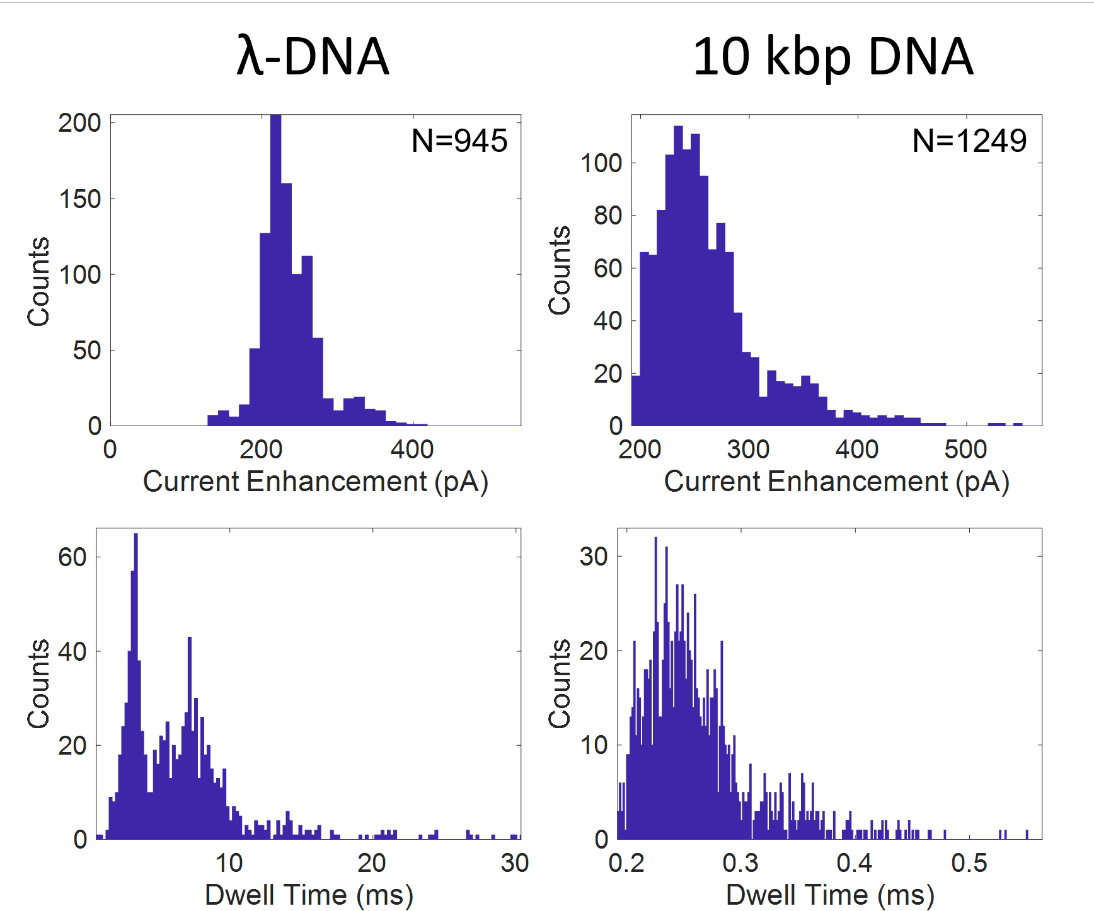

Supplemental Figure 19. Event properties (current enhancement and dwell time) for λ-DNA and 10 kbp DNA at -400 mV applied voltage bias and asymmetric salt conditions (1 M KCl inside the pipette and 4 M KCl outside the pipette). Both KCl solutions were buffered with Tris-EDTA buffer to pH 7.4. The concentration of DNA in both experiments was 500 pM.

**XIV. Event decays allude to current deficit and current excess**

If events should have a pulse-like shape, there are two current transients that must be added to the pulsed shape: one at the beginning of the event and one at the end of the event. Both components 1 and 2 sum to equal the conductive event signature. Component 2 demonstrates the non-equilibrium components of the signal. The initial moments of the event show a current transient that counteracts the current enhanced state while the end of the event extends the duration of the current enhanced state. A mathematical model for the current transients were created in MATLAB in which a hypothetical C(t) and V(t) were created. In one simulation, the voltage, V(t), had a pulse-like trend as shown in Supplemental Figure 17c and the capacitance was constant. This resulted in current transients that matched Component 2. The second simulation made the capacitance, C(t), variable and pulse-like. The result also showed current transients similar to Component 2. If the voltage pulse direction, or the capacitance pulse direction, were switched, then the current transients would be in the opposite direction.

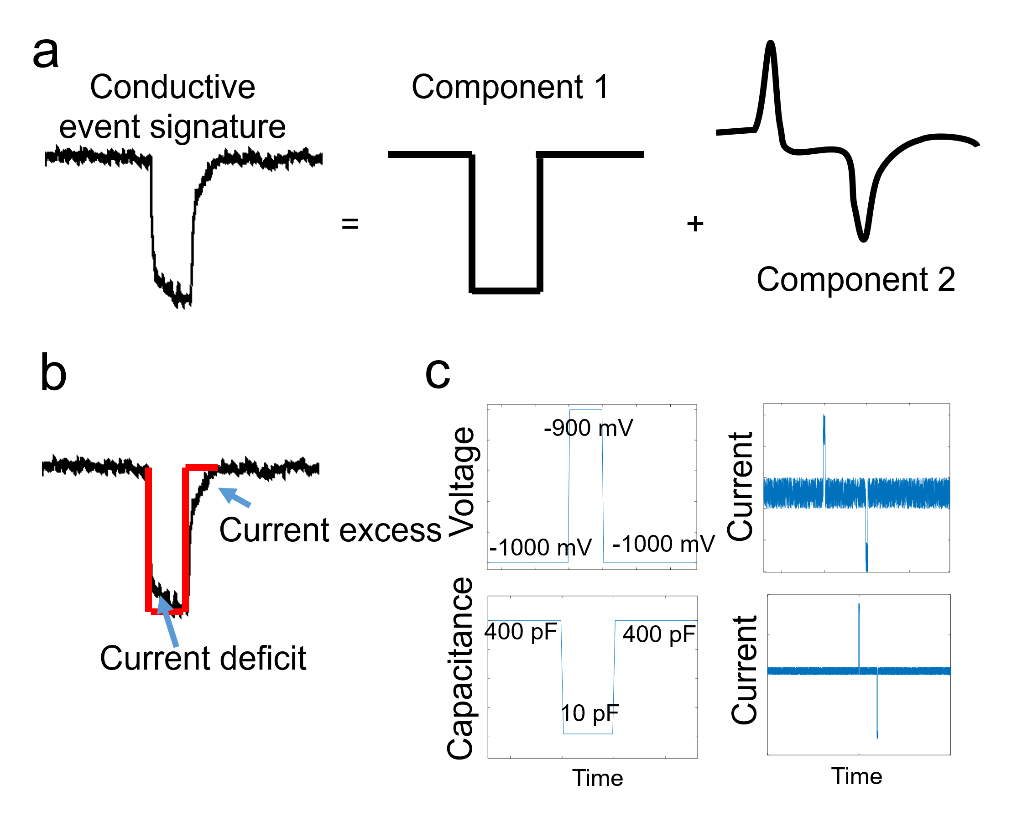

Supplemental Figure 20. (a) The conductive pulse event signature can be described as being composed of a pulse-like component and a current transient component. (b) The leading and falling edge of the event signature show both a current deficit as well as a current excess which prolongs the current enhancement. (c) MATLAB simulation of a variable voltage across a capacitor (top), and a variable capacitance (bottom). Since the cause of the pulse (component 1) is undisputed, only the capacitive component is modelled in order to explain the positive and negative contributions to current at the beginning and end of an event. Random noise was added to simulate a real current measurement setup. The equation used for the simulations was $I=C\frac{dV}{dt}+V\frac{dC}{dt}$. The default values for C and V were 10 pF and -1000 mV.

**XV.** **Fitting exponential decays to events**

The time constants associated with a voltage switch as well as the DNA events were fitted by a single exponential of the form: $\Delta I=Ce^{-rt}$, where C and r are fitting constants. If the exponential shape of the current transient was anything other than a positive current decaying towards zero, the signals were flipped (multiplied by -1) and the equilibrium current value was subtracted. All fits were checked for goodness of fit (R^2^>0.95). For normal capacitance, the exponential decays found at negative voltages simply were multiplied by -1 and could be fitted by an exponential (after equilibrium current was subtracted). If negative capacitance was present, the signals already had a decaying trend and thus only the equilibrium value was subtracted. If the trace was not multiplied by -1, then the decay rate was multiplied by -1 to indicate the presence of negative capacitance. Decay rates that were negative, after this procedure, represented negative capacitance. Unless otherwise noted, the first 20 ms of data (post voltage switch) were used for fitting. For fitting of DNA rise times (i.e., the beginning of the event), the initial data point was the first FWHM point. The subsequent 0.8 ms of the event was then fitted to the exponential ($\Delta I=Ce^{-rt})$ after subtracting the equilibrium value such that the current decayed to zero. Prior to this, only events >1 ms were used to ensure that 0.8 ms of data was obtainable for accurate fitting. For fitting the DNA fall times (i.e., the end of the event), the second FWHM point was used as the initial data point. The subsequent 0.8 ms of data was then used for fitting. For DNA event fitting, no sign (+/-) was given to the decay rate since it was unclear whether the decay represented negative capacitance; specifically as it was defined earlier.

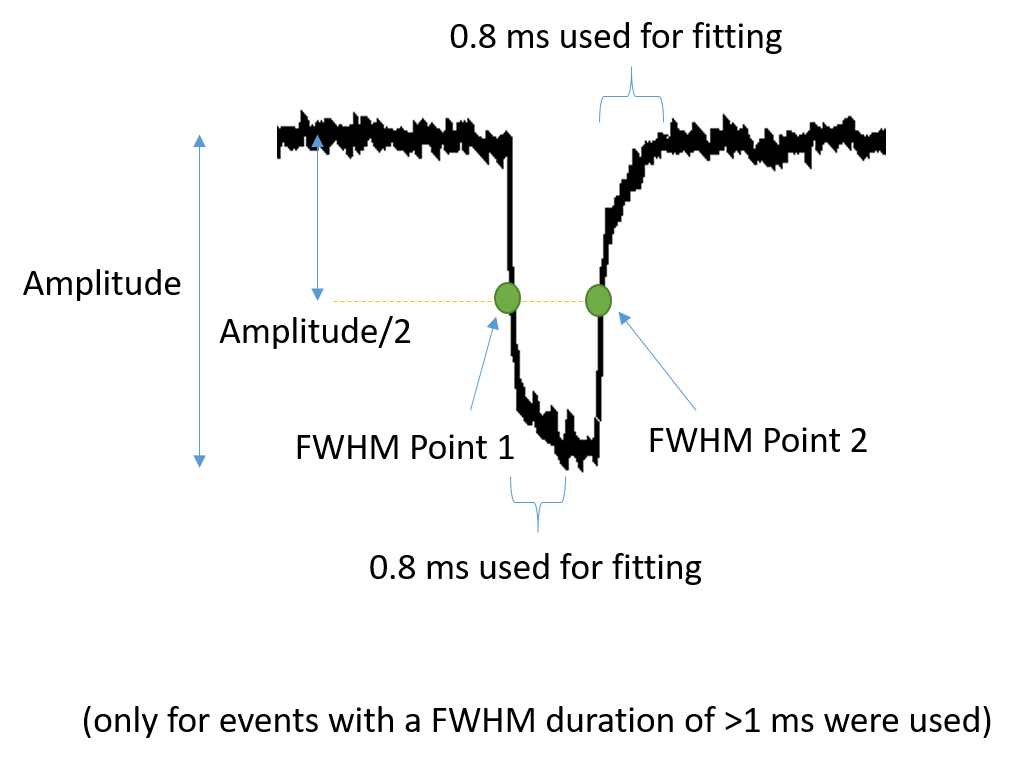

Supplemental Figure 21. Schematic of how the event decay rates were found using the full width half max (FWHM) of each event. Briefly, 0.8 ms of data after each FWHM point was used for fitting an exponential. The decay rate during the initial moments of the event were called the event rise (DNA entry) and the fall back towards the open pore current was termed the event fall (DNA exit).

**XVI. Warburg impedance model**

The Warburg impedance model is commonly used in electrical impedance spectroscopy (EIS) for describing electrochemical systems. The Warburg impedance is especially useful for modeling diffusion processes within electrochemical systems and can further be decomposed into an infinite series of parallel RC circuits. The Warburg impedance acts both as a resistor and a capacitor since it induces a magnitude and phase shift between the voltage and current of the circuit that varies with frequency^3^. The two Warburg elements, R_W_ and C_W_, correspond to the resistance to concentration polarization and the activation energy for polarization (i.e., charging process). By analyzing the voltage drop across each element, it becomes clear that the applied voltage is spread across the pore and the taper. The voltage across the pore is reduced by concentration polarization which could fluctuate throughout an experiment depending on the flux of ions traveling through the pore. Since the diffusion process would likely be altered by the presence of DNA within the taper length of the nanopipette, the inclusion of a Warburg element also explains why the effective capacitance was different for the rising and falling edges of the event signature at low salt (10 mM KCl).
